# Systematic Analysis of GARP- and EARP-Mediated Trafficking Pathways Using Lysosomal Proteomics

**DOI:** 10.64898/2026.10.08.757561

**Authors:** Bhavya Sivalingam, Janathan Altuzar, Stefan Walter, Joost C. M. Holthuis, Bianca M. Esch, Florian Fröhlich

**Author notes:** Corresponding authors: (F.F.); (B.M.E).

## Abstract

Eukaryotic cells utilize tethering complexes for sorting of proteins within the endo-lysosomal system. The Golgi-associated retrograde protein (GARP) complex and the endosome-associated recycling protein (EARP) are two tethering complexes involved in endosomal recycling pathways. Both complexes share three common subunits (VPS51, VPS52, and VPS53), with each relying on one specific subunit: VPS54 for GARP and VPS50 for EARP. Mutations in both complexes are linked to various neurodegenerative disorders, such as progressive cerebello-cerebral atrophy. While model cargoes have been described, the cargo spectrum of both complexes remains largely elusive. Here, we combined lysosomal proteomics with knockdowns of GARP and EARP to analyze their cargo spectra. We first generated a deep lysosomal reference proteome. Using the lysosome as a terminal reference point for mis-sorted proteins, we show that GARP depletion causes a loss in lysosomal hydrolases and accumulation of non-lysosomal cargo. EARP depletion results in lysosomal accumulation of secreted proteins. Our findings serve as a reference to study GARP and EARP complexes and provide a comprehensive overview of the lysosomal proteome. We demonstrate that a systematic phenotypic analysis of trafficking mutants is necessary to understand the pathological consequences of their dysfunction.

**Statement of significance of the study:** This study provides a systematic analysis of the lysosomal proteome using mass spectrometry-based proteomics. In comparison to other studies, our work represents one of the most comprehensive analyses available to date. By combining this method with knockdowns of the GARP and EARP complexes, we establish the lysosomal proteome as a reference for identifying mis-sorted proteins. This approach enables us to characterize the global effects of GARP and EARP depletion. Our findings provide new insights into the cargo spectra of these trafficking complexes and establish a methodological framework for investigating recycling defects and associated protein mis-sorting.

## Introduction

The sorting of proteins retrieved from endocytosis is an essential process to maintain cellular homeostasis. After the scission of endocytic vesicles from the plasma membrane, these carriers fuse with early endosomes. Subsequently, the fate of the endocytosed receptors is determined by the endosomal sorting machinery. These proteins can be incorporated into intraluminal vesicles, leading to their transport to the lysosome, where they are degraded (Malerød et al., 2011; Raiborg and Stenmark, 2009; Wenzel et al., 2018; Wollert and Hurley, 2010). Proteins can also be recycled to the plasma membrane in tubular structures emerging from recycling endosomes or are transported to other compartments such as the Golgi apparatus (Bonifacino and Rojas, 2006; Simonetti et al., 2023; Toshima and Toshima, 2024).

Endosomal sorting depends on multiple factors such as the sorting nexins (Carlton et al., 2005, 2005; Cullen, 2008), the retromer complex (Arighi et al., 2004; Seaman, 2004; Seaman et al., 1998), SNARE proteins/complexes (Hong and Lev, 2014; Kümmel and Ungermann, 2014; Söllner et al., 1993), and various tethering complexes required for the capturing of tubular-vesicular carriers. Two important tethering complexes in the endo-lysosomal system are the Golgi-associated retrograde protein (GARP) complex and the endosome-associated recycling protein (EARP) complex (Liewen et al., 2005; Schindler et al., 2015). GARP and EARP belong to the family of complexes associated with tethering containing helical rods (CATCHR). Both complexes share the three subunits VPS51/Ang2, VPS52, and VPS53 and each harbors a specific subunit, either VPS54 (GARP) or VPS50/Syndetin (EARP).

The role of both complexes has been described in detail for several model cargoes such as the transferrin receptor, or the mannose-6-phosphate receptor (Pérez-Victoria et al., 2008). In addition, mis-function of subunits of both complexes have been linked to various phenotypes such as disrupted lipid homeostasis (Eising et al., 2019; Fröhlich et al., 2015; Wei et al., 2017), impaired Golgi function resulting in impaired protein glycosylation (Khakurel et al., 2025, 2022, 2021), and disrupted intracellular trafficking (Hirata et al., 2015; Pérez-Victoria et al., 2008). These phenotypes are associated with neurological developmental disorders and neurodegenerative diseases (Beilina et al., 2020; Gershlick et al., 2019; Ghosh et al., 2026). Mutations of the GARP-specific VPS54 subunit are also linked to neurodegenerative disorders. For example, a point mutation in VPS54 on mice leads to the wobbler mouse phenotype with progressive upper and lower motor neuron degeneration, neuromuscular weakness, and muscle wasting (Pérez-Victoria et al., 2010; Petit et al., 2020). This mouse model is widely used for studying amyotrophic lateral sclerosis (ALS) (Duchen and Strich, 1968; Schmitt-John et al., 2005). A systematic proteomic analysis of wobbler mouse tissue revealed the mis-sorting of various proteins at the Golgi apparatus. Despite their severe pathological impact, a comprehensive analysis of the consequences of mutations in GARP and EARP complexes on endo-lysosomal trafficking of protein cargo in higher eukaryotic systems is lacking.

A common phenotype of GARP- and EARP misfunction is the accumulation of recycling cargo in lysosomes. Although, lysosomal proteins are usually low abundant in proteomics analyses, lysosomes are one of the most studied organelles in enrichment studies. Protocols for lysosomal immunoprecipitation have been developed and used for the characterization of lysosomal disorders (Abu-Remaileh et al., 2017; Saarela et al., 2024). Thus, the proteomic analysis of purified lysosomes from GARP- and EARP- mutant cells should allow the determination of missorted cargo proteins and allow a systematic analysis of their cargo spectrum. A recently published study examined the molecular environment of GARP and EARP complexes, employing a complementary approach to investigate their cargo spectrum (Aragon-Ramirez et al., 2026).

Here, we developed a lysosomal purification protocol based on the immuno-isolation of LAMP1-GFP from HeLa cells. We combined this approach with siRNA-mediated knockdown of the mannose-6-phosphate receptors to test the feasibility of the method, and to determine changes in the lysosomal proteome. While the observed changes are modest, these experiments leave little room for adaptive processes occurring, for instance in knockout cells. Therefore, our approach provides a realistic picture of the cargo spectrum of the receptors. After carefully determining the lysosomal reference proteome in HeLa cells, we analyzed the knockdowns of all GARP and EARP subunits and determine their effects on the lysosomal proteome. The GARP knockdown led to a reduction in lysosomal hydrolases and lipid transport proteins, while the EARP knockdown resulted in an increase in extracellular and secreted proteins in the lysosome. The VPS53KD had the most severe effects and exhibited a combined phenotype, while the VPS52KD and VPS51KD showed the same trends but with decreasing severity. We also confirm several of our proteomic findings by imaging of newly identified cargo proteins and compare our results with previous studies of the full proteome and the Golgi proteome of wobbler knockout mice. In summary, we present an in-depth lysosomal proteome in combination with siRNA knockdowns to determine the individual cargo spectra of the GARP and EARP tethering complexes.

## Results

### Establishment of a robust lysosomal enrichment strategy

To characterize the effects of GARP and EARP depletions on the lysosomal proteome, we aimed at establishing a protocol allowing the isolation of intact lysosomes under conditions where the complexes are acutely depleted. Various protocols for lysosomal purifications have been developed relying on immuno-isolation using lysosomal marker proteins or the labeling of lysosomes with supra-magnetic particles (Akter et al., 2023; Tharkeshwar et al., 2020). We have recently used a purification strategy based on a GFP-tagged version of the LAMP1 protein that is only minimally overexpressed in HeLa cells (Fig. 1a) (Adeosun et al., 2026). First, we wanted to characterize this purification method in comparison to other lysosomal purification strategies and to establish the pulldowns in siRNA treated cell lines.

**Fig. 1:**
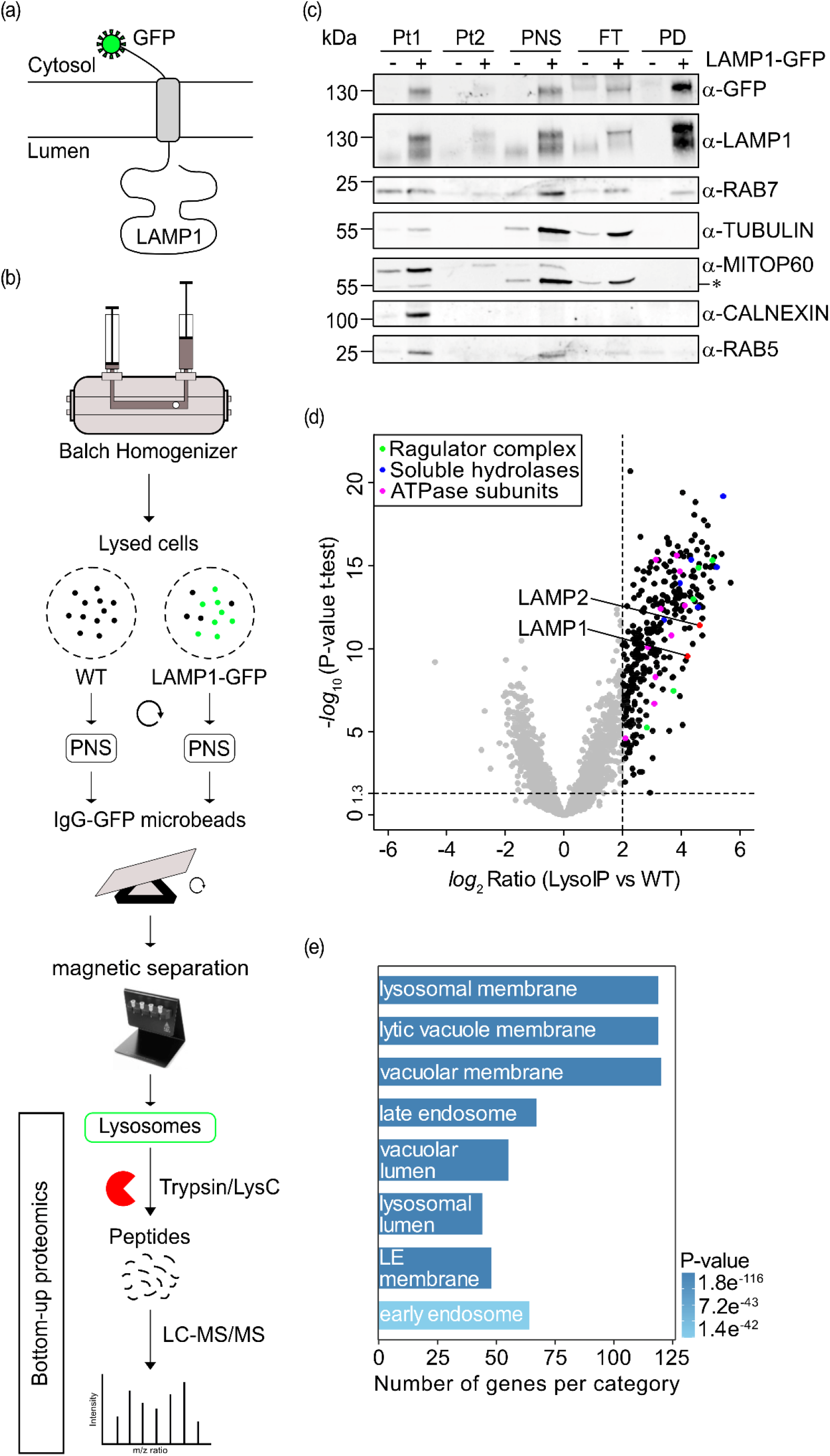
Immunoprecipitation of intact lysosomes via a GFP-tagged LAMP1 construct. Schematic representation of (a) the LAMP1-GFP construct stably expressed in HeLa cells, and (b) the organellar pulldown of the lysosomes using the GFP magnetic beads and their subsequent sample preparation for mass spectrometric analysis. (c) 2.5 v/v % of the nuclear debris pellet (Pt1), light membrane fraction pellet (Pt2), post-nuclear supernatant (PNS), flow-through from the magnetic column (FT), and 12.5 v/v % of the eluate bound to the GFP beads (PD) from HeLa cells with and without the LAMP1-GFP construct on the western blot decorated with antibodies against GFP (construct), LAMP1 (lysosomes), RAB7 (late endosomes), TUBULIN (cytosol), MitoP60 (mitochondria), CALNEXIN (ER), and RAB5 (early endosomes). *Residual α-TUBULIN signal following membrane stripping. (d) Volcano plot of lysosomal immunoprecipitation (Lyso-IP) using LAMP1-GFP compared to wild-type (WT) cells highlighting known lysosomal proteins LAMP1 and LAMP2 in red, ATPase subunits including VA0D1, VATG1, VATB2, VATC1, VATE1, VAS1, VATF, VPP1, VATH, and VATD in magenta, Ragulator complex subunits (LTOR1-5) in green, and soluble lysosomal hydrolases such as CATD, CATB, CATZ, HEXA, HEXB, and BGLR in blue. Proteins with *log*_2_ Ratio > 2 and -*log*_10_ P-value > 1.3 are considered enriched in this study (n=15). (e) The GO term analysis of the enriched proteins shows the top GO terms as lysosomal membrane or lysosome-associated proteins.

The purification strategy was based on the mild lysis of cells, the immuno-isolation of lysosomes with magnetic anti-GFP beads, followed by elution and preparation for mass spectrometry-based proteomics (Fig. 1b). Western blot analysis of the purified lysosomes revealed a good enrichment of both, endogenous LAMP1 and GFP-tagged LAMP1, as well as the late endosomal/lysosomal RAB GTPase Rab7 (Fig. 1c). While we also observed a mild enrichment of the early endosomal RAB GTPase Rab5, other organelles such as the mitochondria (marked by MitoP60), the ER (marked by calnexin) or the tubulin network (marked by tubulin) were not in enriched in our purifications (Fig. 1c). Based on the observed lysosomal enrichment, we subjected our samples to protein digestion, followed by mass spectrometry to determine the lysosomal proteome. Therefore, we compared the enriched lysosomal purification of LAMP1-GFP cells (LysoIP) to a mock purification from cells only expressing the endogenous LAMP1 gene (WT) (Fig. 1d). As expected, this analysis showed enrichment of known lysosomal proteins such as LAMP1 and LAMP2, ATPase subunits (VA0D1, VATG1, VATB2, VATC1, VATE1, VAS1, VATF, VPP1, VATH, and VATD), Ragulator complex subunits (LTOR1-5), and soluble lysosomal hydrolases (CATD, CATB, CATZ, HEXA, HEXB, and BGLR) demonstrating the enrichment of intact lysosomes. To confirm the lysosomal enrichment, we performed GO term analyses of the at least two-fold enriched proteins in the LAMP1-GFP purifications. This analysis revealed a strong over-representation of proteins related to lysosomes, lysosomal lumen and also including GO terms such as late endosome (Fig. 1e). Similarly, comparing the enriched lysosomal fraction directly to a post-nuclear supernatant revealed a strong enrichment of lysosomal proteins (Fig. S1a, b). Taken together, our results suggest a highly significant enrichment of intact lysosomes from cells only minimally overexpressing LAMP1-GFP.

### Benchmarking the lysosomal proteome

Next, we compared our lysosome purifications to two recent benchmark lysosomal proteomics studies (Akter et al., 2023; Hein et al., 2025). First, we subdivided the enriched protein fractions into bins and analyzed the enrichment of GO terms within each bin, similar to previous analyses of yeast vacuoles (Eising et al., 2019) (Fig. 2a). All bins with a fold enrichment greater than two showed only GO terms related to lysosomal compartments (Fig. 2a, b). The bin with one- to two-fold enriched proteins still contained lysosomal GO terms (vacuolar membrane, lysosomal membrane, lytic vacuole membrane) but also other GO terms, such as focal adhesion and cell-substrate junction (Fig. 2a, b). To use a rather stringent cutoff, we considered all proteins with a fold change larger than 2 as enriched in our lysosomal fraction. This analysis yielded a list of 265 proteins that we consider to represent the lysosomal proteome. We compared this fraction to the other two lysosomal proteome datasets as a reference. The comparison revealed a high overlap between ours and the other studies, with 120 proteins identified in our data and at least one of the other two studies (Fig. 2c). Additionally, we identified 145 proteins as lysosomal-enriched that were not previously annotated in either of the other two datasets. GO term analysis of the 145 proteins exclusively enriched in our dataset again showed mainly lysosome-related GO terms, further supporting the validity of the chosen cut-off values (Fig. 2d). Next, we highlighted the proteins identified either in Akter et al., 2023 (magenta) or Hein et al., 2025 (blue, black for identification in both datasets) in our dataset (Fig. 2e). This plot revealed a high overlap of the compared datasets with only a smaller fraction of proteins that were identified as lysosomal proteins only in the other studies. Highlighting the proteins which were identified in our lysosomal enrichment and in at least one other dataset (red = Akter et al., 2023, or Hein et al., 2025; black = identified in all three datasets) confirms the strong enrichment of intact lysosomes in our study (Fig. 2f). Only 25 proteins which were identified in Akter et al., 2023 were not detected in our dataset. From lysosomal proteins identified in Hein et al., 2025, 89 proteins were not detected in our dataset. In summary, by using a stringent statistical cutoff we generated a high-confidence lysosomal proteome data set for HeLa cells, identifying 265 lysosomal proteins.

**Fig. 2:**
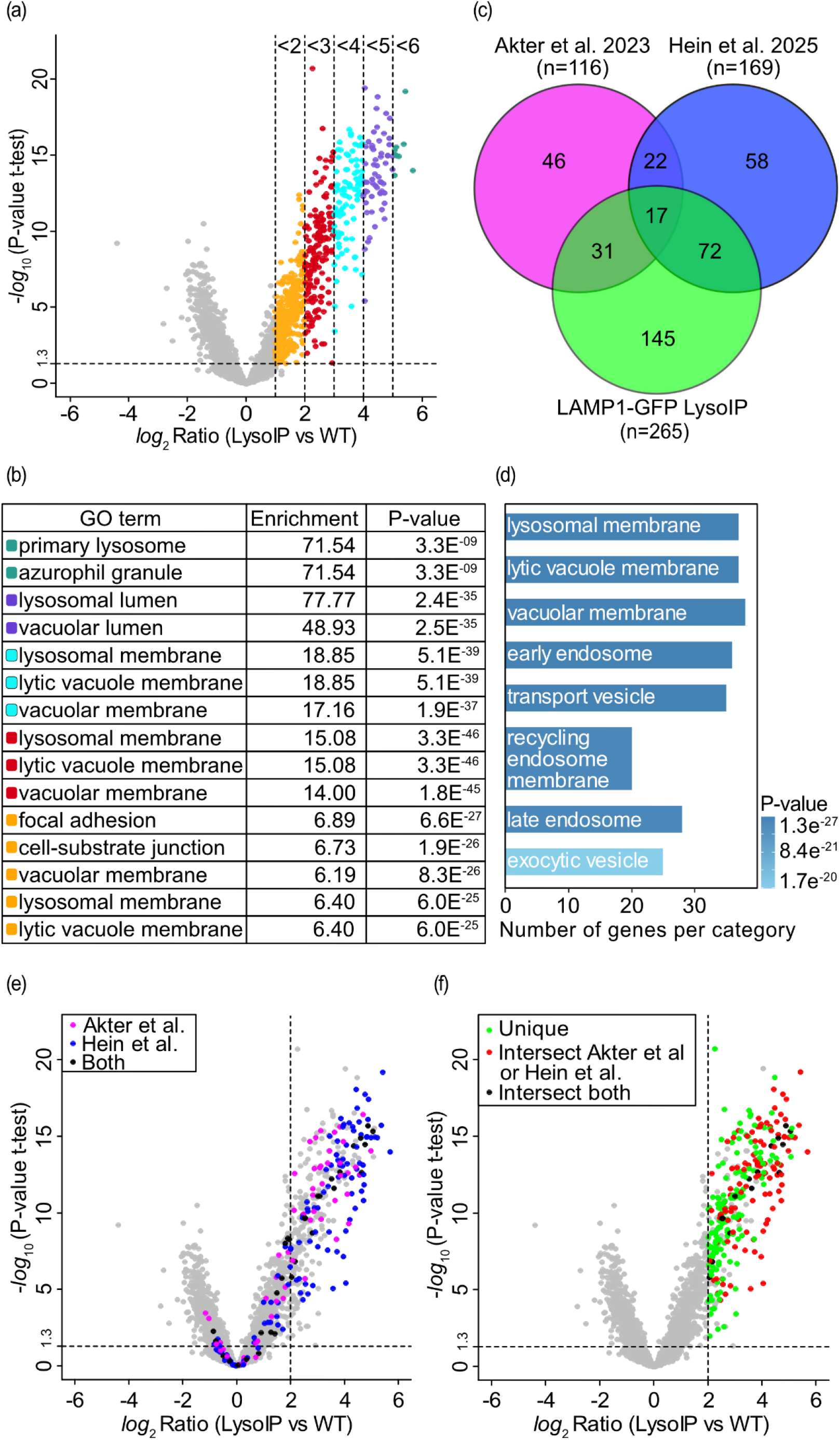
Lyso-IP enrichment analysis highlighting unique lysosomal proteins. Analysis of the Lyso-IP enrichment by (a) binning the intervals of the *log*_2_ Ratio, and (b) identifying the top GO terms for each interval. Enrichment with a *log*_2_ Ratio Lyso-IP > 2 corresponds to lysosomal GO terms. (c) Venn diagram comparing enriched proteins (*log*_2_ Ratio Lyso-IP > 2, -*log*_10_ P-value > 1.3, n = 265) with curated lysosomal protein lists from Akter et al. 2023 (n = 116) and Hein et al. 2025 (n = 169). Among the enriched proteins, 48 identified lysosomal proteins overlap with lysosomal proteins from Akter et al., 2023. 89 identified lysosomal proteins overlap with Hein et al., 2025. 17 identified lysosomal proteins overlap with both datasets (Akter et al., 2023; Hein et al., 2025). In total, 120 lysosomal proteins from our dataset were previously annotated lysosomal proteins. (d) GO term enrichment of the remaining 145 unique proteins reveals lysosome-related terms as the most significantly enriched. (e) Lyso-IP scatterplot depicting enriched lysosomal proteins reported in Akter et al., 2023 (magenta), Hein et al., 2025 (blue), or both (black). From Akter et al., 2026 (116 lysosomal proteins annotated), we detected 91 proteins (25 proteins undetected), among which 48 proteins were lysosomal enriched in our dataset (overlap seen in Venn diagram) and 43 proteins were below the threshold. From Hein et al., 2025 (169 lysosomal proteins annotated), we detected 145 proteins (24 proteins undetected), among which 89 proteins were lysosomal enriched in our dataset (overlap seen in Venn diagram), and 56 proteins were not enriched in our dataset. (f) Lyso-IP scatterplot showing uniquely enriched proteins from our dataset (green), proteins overlapping with the lysosomal dataset of Akter et al. (red) or Hein et al. (red), and proteins overlapping with both datasets (black).

### Determination of the lysosomal proteome following siRNA-mediated depletion of transport receptors

Having established a lysosomal reference proteome for HeLa cells, we next tested whether siRNA-mediated knockdown of trafficking receptors leads to detectable changes in the proteome of immuno-isolated lysosomes. Although siRNA-mediated interference with gene expression overall produces milder phenotypes in comparison to gene KO strategies, a clear advantage of the KD approach is that they leave cells less time to adapt to a loss of protein function. As proof-of-principle, we analyzed the impact of siRNA-mediated downregulation of the well-characterized mannose-6-phosphate receptor (M6PR) on the sorting of lysosomal cargo (Fig. 3a). HeLa cells express both a cation-dependent M6PR and a cation-independent M6PR (IGF2R). Both receptors can compensate for the loss of each other (Gauthier et al., 2024; Sleat and Lobel, 1997). Therefore, we performed a double siRNA-mediated knockdown of M6PR and IGF2R. Depletion of M6PR was confirmed by Western blotting (Fig. S1c), while we lacked a functional antibody for IGF2R. However, knockdown of IGF2R was confirmed by proteomics (Fig. 3d).

**Fig. 3:**
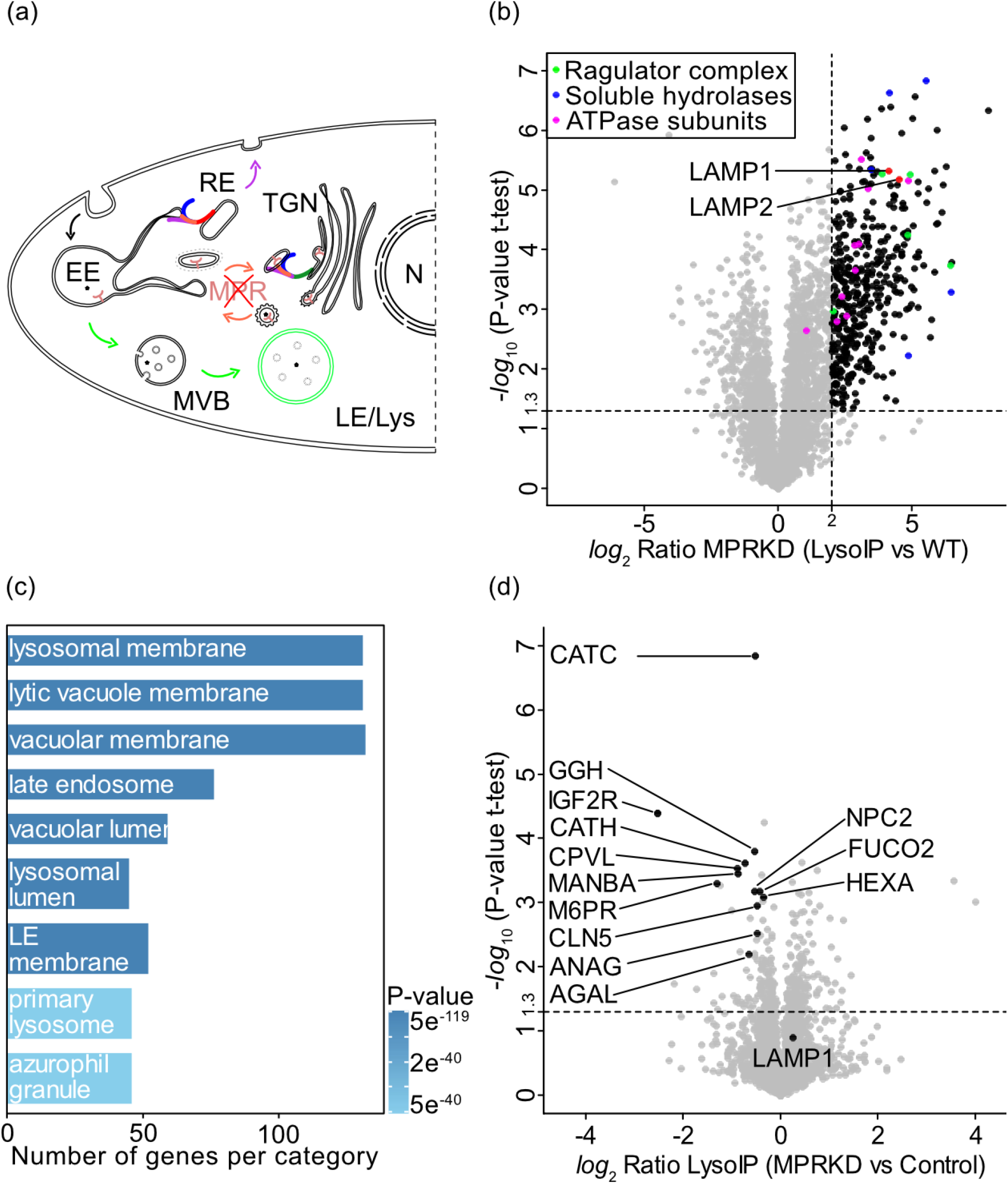
Impairment of lysosomal enzyme targeting upon MPR knockdown. (a) Schematic overview of endocytic trafficking pathways showing that simultaneous knockdown of cation-dependent and cation-independent mannose-6-phosphate receptors (MPRs) impairs transport of lysosomal enzymes from the trans-Golgi network (TGN) to early endosomes (EE), which mature into late endosomes (LE) and subsequently fuse with lysosomes (Lys). (b) Scatterplot of Lyso-IP from MPR knockdown cells showing enriched proteins (*log*_2_ Ratio Lyso-IP > 2, -*log*_10_ P-value > 1.3) in black. Known lysosomal proteins LAMP1 and LAMP2 in red, ATPase subunits including VA0D1, VATG1, VATB2, VATC1, VATE1, VAS1, VATF, VPP1, VATH, and VATD in magenta, Ragulator complex subunits (LTOR1-5) in green, and soluble lysosomal hydrolases such as CATD, CATB, CATZ, HEXA, HEXB, and BGLR in blue are highlighted. (c) GO term analysis of the enriched proteins reveals lysosome-associated proteins as the most significantly enriched proteins (n=3). (d) Volcano plot depicting the fold-change of proteins upon MPR knockdown in comparison to the control Lyso-IP. Previously known MPR cargoes such as NPC2, HEXA, CATH, CATC, GGH, CPVL, MANBA, CLN5, ANAG, AGAL, and FUCO2 show decreased enrichment in the MPR depleted lysosomes (n=3).

greater than 2 were again lysosomal proteins, as shown by GO term analysis (Fig. 3b, c). Comparison of the lysosomal enrichments from Lyso-IPs from LAMP1-GFP expressing cells either treated with control siRNA or M6PR and IGF2R siRNA revealed differences in the lysosomal content. While the changes were subtle (Fig. 3d; fold change of around 1), lysosomal proteins that were downregulated in cells with knocked-down M6PRs were luminal lysosomal proteins (including NPC2, HEXA, CATH, CATC, GGH, CPVL, MANBA, CLN5, ANAG, AGAL, and FUCO2) and have been previously shown to be cargo of the M6PR pathway (Fig. 3d) (Sleat et al., 2006; Willenborg et al., 2005). This approach allowed us to maintain relatively stable lysosomal protein levels, avoiding the large-scale proteomic shifts associated with complete gene knockouts, which often trigger compensatory metabolic adaptations. Together, our results show that siRNA knockdowns can be used to investigated the cargo spectrum of trafficking enzymes, with the lysosomal proteome serving as a read-out.

### Systematic analysis of GARP- and EARP-mediated trafficking

After confirming that Lyso-IPs combined with siRNA-mediated knockdowns of cargo receptors can reveal cargos by investigating the lysosomal proteome, we next characterized knockdowns of the GARP and EARP complexes. GARP is involved in the recycling of proteins from endosomes to the Golgi apparatus, while EARP is described to be involved in the recycling of proteins from endosomes to the plasma membrane (Fig. 4a) (Pérez-Victoria et al., 2008; Schindler et al., 2015). Importantly, both complexes share the subunits VPS51, VPS52, and VPS53. The protein VPS54 is specific to the GARP complex, in contrast to the VPS50 protein, which is an EARP-specific subunit (Fig. 4b). Knockdowns of VPS50, VPS51, VPS53, and VPS54 were first confirmed by western blotting (Fig. S1c). While the knockdowns had varying efficiencies, imaging of lysosomes showed the expected phenotypes of enlarged perinuclear lysosomes in VPS50, VPS53 and VPS54 knockdown cells (Fig. S1d) (Feinstein et al., 2014; Fröhlich et al., 2015; Pérez-Victoria et al., 2008; Wei et al., 2017). Triplicates of knockdowns with control siRNA, VPS50-, VPS51-, VPS52-, VPS53-, and VPS54-siRNA in LAMP1-GFP expressing cells were used for immuno-isolation of lysosomes. The resulting lysosomal enrichments were analyzed by mass spectrometry-based proteomics and compared to immunoprecipitations of cells expressing the untagged wild-type copy of LAMP1 and the corresponding siRNA (Fig. S2). All previously defined lysosomal proteins are highlighted in green, showing that the overall lysosomal purifications also worked in cells treated with siRNAs. To analyze the effect of knockdowns of GARP and EARP subunits, we first performed a soft clustering (Kumar and Futschik, 2007) of all proteins which were significantly enriched (fold change greater than 2) in one of the Lyso-IPs (Fig. 4c). The soft clustering revealed six different protein clusters. Two of the clusters (3 and 6) showed very specific phenotypes for proteins enriched or depleted in VPS50 KD cells. In contrast, the knockdowns of VPS52 and VPS53 showed mostly similar results to the knockdown of VPS54, suggesting that the effects of GARP KDs were dominant over those from the EARP complex. Knockdown of the VPS51 subunit showed the weakest effects, also in line with a previously observed minor role in complex formation (Conibear et al., 2003; Gershlick et al., 2019; Luo et al., 2011; Siniossoglou and Pelham, 2002). The results from the soft-clustering were confirmed by hierarchical clustering (Fig. 4d). The analysis shows that each replicate of the knockdowns is most similar to each other. In addition, the large hierarchical distance between the VPS50KD and VPS54KD samples demonstrated the differences between a GARP and EARP-specific knockdown, with the common subunits clustering in between (Fig. 4d).

**Fig. 4:**
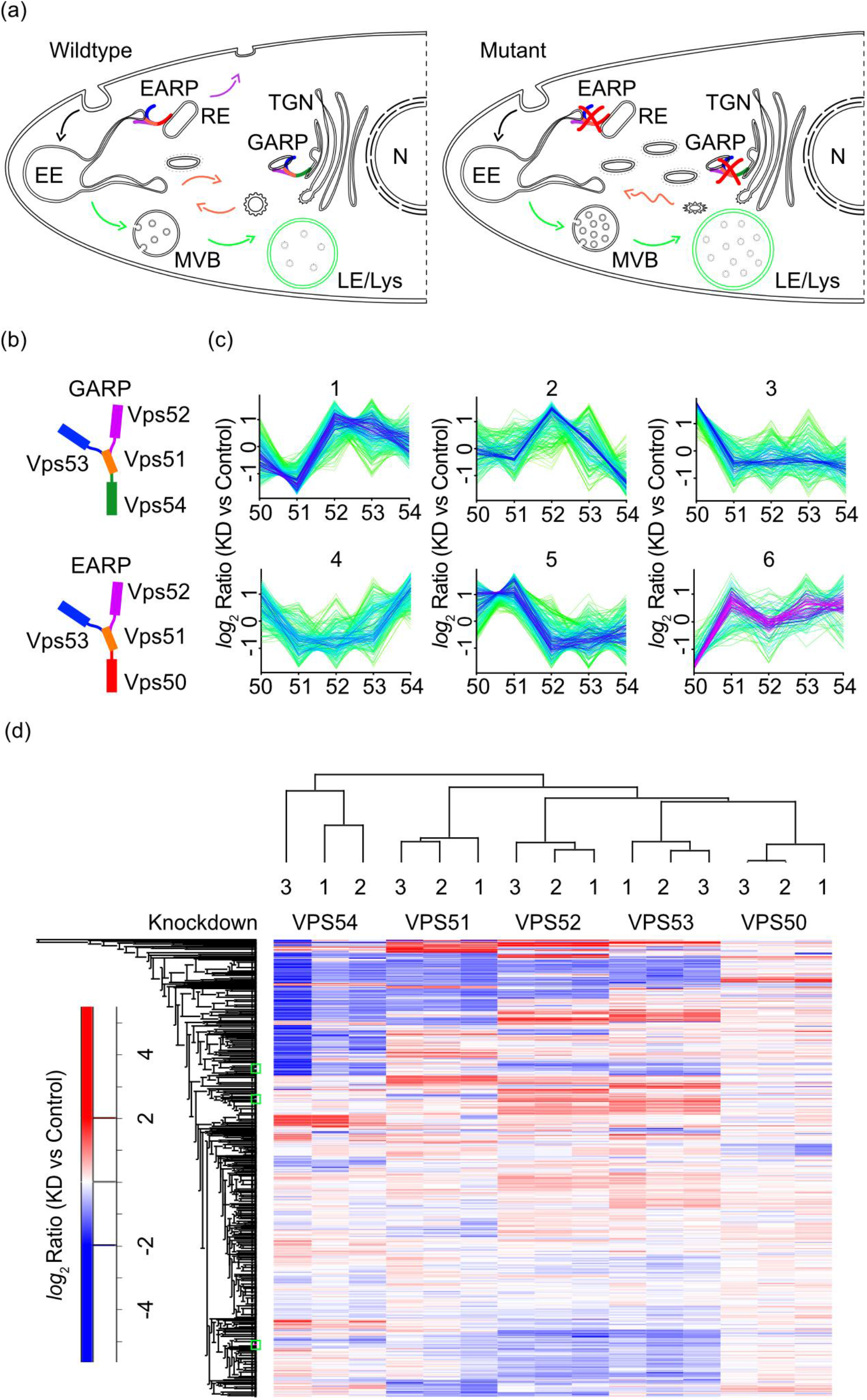
Lysosomal proteome profiling following GARP/EARP complex knockdowns. (a) Working model illustrating the roles of the GARP and EARP complexes in endocytic trafficking pathways under wild-type conditions (left) and the proposed effects upon GARP/EARP knockdown (right). In wild-type cells, the EARP complex on recycling endosomes (RE) tethers endosome-derived vesicles (carrying cargoes or receptors for exocytosis), facilitating their pinch-off from early endosomes (EE) and subsequent fusion with the RE via SNARE proteins. The GARP complex on the trans-Golgi network (TGN) tethers retromer-associated vesicles (containing cargoes or receptors to be recycled back to the Golgi) and enables their fusion with the TGN via SNAREs. We propose that, upon GARP/EARP knockdown, retromer-associated vesicles fail to fuse with the TGN, and endosome-derived vesicles fail to pinch off from the EE. (b) Cartoon representation of the GARP and EARP complexes, highlighting their shared subunits (Vps51, Vps52, Vps53) and their unique subunits (Vps54 for GARP, and Vps50 for EARP). (c) Soft clustering of the enriched proteins (*log*_2_ Ratio Lyso-IP > 2, -*log*_10_ P-value > 1.3) in Lyso-IPs from GARP/EARP knockdowns to profile proteomic changes of the lysosomes. (d) Clustering analysis using a heatmap to highlight changes of enriched proteins in Lyso-IP from GARP/EARP knockdowns relative to control, illustrating alterations in the lysosomal proteome. Proteins de-enriched in knockdown lysosomes relative to control are shown in blue, while proteins enriched in knockdown lysosomes are shown in red, with gradient colors reflecting fold-change magnitude.

### Disruption of lysosomal lipid homeostasis after combined depletion of GARP and EARP complexes

To further investigate the effects of GARP and EARP depletion on the lysosomal proteome, we compared the VPS53KD Lyso-IP to a control Lyso-IP with scrambled siRNA. VPS53-depleted cells showed a de-enrichment of 43 enzymes which were identified as mainly lysosomal (lysosomal lumen, vacuolar lumen, lysosomal membrane, lytic vacuole membrane, vacuolar membrane), but also as early endosomal by GO term analysis (Fig. 5a, b). Lysosomal proteins which were significantly de-enriched included the carboxypeptidase CPVL, the fucosidase FUCO, the ESCRT-III component CHMP3, pro-cathepsin H (CATH), the palmitoyl-protein thioesterase 1 (PPT1), and the alpha-N-acetylgalactosaminidase (NAGAB). Another de-enriched enzyme in the VPS53KD is the lysosomal cholesterol transporter NPC2 (Fig. 5a). GARP misfunction was previously linked to impairments in cholesterol trafficking (Wei et al., 2017). In addition, other lysosomal lipases such as the lysosomal acid lipase (LICH), glucosylceramidase (GBA1), and phospholipase PAG15 were similarly depleted from lysosomes, highlighting a possible disruption of lysosomal lipid homeostasis in VPS53KD cells. These findings indicate an impaired delivery of lysosomal proteins without GARP- and EARP-mediated trafficking. We hypothesized that these proteins could be misrouted to the Golgi apparatus under these conditions. To test this, we correlated the VPS53-depleted lysosomal proteome to the Golgi proteome from VPS54-deficient wobbler mice (Fig. S3a) (Petit et al., 2020). This analysis revealed a subset of proteins which is deficient in VPS53KD lysosomes and enriched in the Golgi of wobbler mice such as PPT1, NAGAB, NPC2, glucuronidase (BGLR), peptidase DPP2, galactosidase (AGAL), FUCO, and acetylchitobiase DIAC (Fig. S3a).

**Fig. 5:**
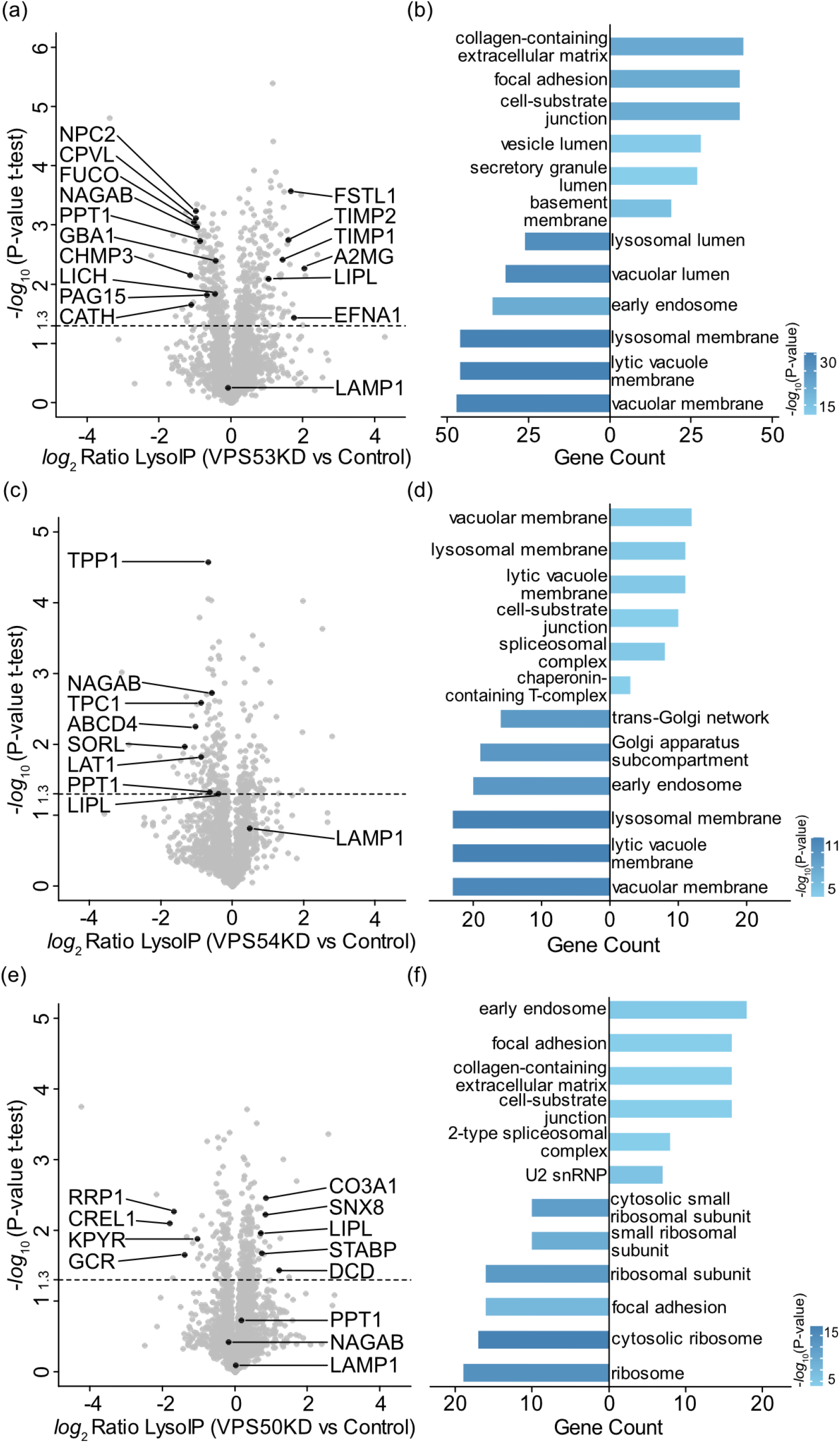
GARP/EARP knockdown impairs lysosomal enzyme targeting and promotes cargo retention. (a) Volcano plot of Lyso-IP from VPS53 (GARP and EARP complex) knockdown cells showing de-enrichment of lysosomal enzymes including carboxypeptidase CPVL, the fucosidase FUCO, the ESCRT-III component CHMP3, pro-cathepsin H (CATH), the palmitoyl-protein thioesterase 1 (PPT1), alpha-N-acetylgalactosaminidase (NAGAB), lysosomal cholesterol transporter NPC2, lysosomal acid lipase (LICH), glucosylceramidase (GBA1), and phospholipase PAG15. Whereas, enrichment of a secreted proteins such as Follistatin-related protein 1 (FSTL1), metalloproteinase inhibitors TIMP1 and TIMP2, alpha-2-macroglobulin (A2MG), lipoprotein lipase (LIPL), and cell surface receptor EFNA1 was observed. (b) GO term analysis of proteins significantly (-*log*_10_ P-value > 1.3) enriched or de-enriched in VPS53 knockdown Lyso-IP. Top enriched terms (right) correspond primarily to extracellular proteins, whereas the top de-enriched terms (left) correspond to lysosomal proteins. (c) Scatterplot of Lyso-IP from VPS54 (GARP-specific) knockdown cells showing de-enrichment of lysosomal enzymes including NAGAB, two pore channel protein 1 (TPC1), lysosomal cobalamin transporter (ABCD4), Sortilin-related receptor (SORL), amino acid transporter (LAT1), lysosomal serine protease TPP1, PPT1, and LIPL. (d) GO term analysis shows that significantly enriched (right) proteins are moderately associated with lysosomal and extracellular terms, whereas significantly de-enriched (left) proteins are strongly linked to lysosomal and Golgi components. (e) VPS50 (EARP-specific) knockdown lysosomes show significant accumulation of the extracellular and endosomal proteins, such as collagen chain CO3A1, sorting nexin SNX8, LIPL, metalloprotease STABP, and dermcidin (DCD), with no significant difference in lysosomal enzymes (NAGAB and PPT1), as illustrated in the volcano plot. (f) GO term analysis of VPS50 knockdown Lyso-IP shows that enriched (right) proteins are primarily associated with extracellular components, while de-enriched (left) proteins are mainly linked to cytosolic components.

At the same time, GO term analysis showed an enrichment of secreted proteins in VPS53KD Lyso-IPs (collagen-containing extracellular matrix, focal adhesion, cell-substrate junction, secretory granule lumen; Fig. 5a, b). Examples are Follistatin-related protein 1 (FSTL1), metalloproteinase inhibitors TIMP1 and TIMP2, alpha-2-macroglobulin (A2MG), lipoprotein lipase (LIPL), and cell surface receptor EFNA1, leading to the hypothesis of impaired recycling of extracellular components without functional GARP and EARP complex. Acute downregulation of the other two common subunits of the GARP and EARP complex demonstrated similar phenotypes as visible in the corresponding GO term analysis (Fig. S4c, d, e, f). De-enriched GO terms were mainly lysosome-related, while enriched proteins were sorted into GO terms corresponding to secreted proteins like focal adhesion or secretory granule lumen. Decreasing numbers of counts for these GO terms from VPS53-KD, to VPS52-KD, to VPS51-KD demonstrated the higher impact of the VPS53 subunit on the lysosomal proteome and is in line with observations from the hierarchical clustering analysis (Fig. 5b and S4d, f). Comparison of these findings to the control knockdowns with the M6PR receptors (Fig. S4a, b) demonstrated the specificity of the observed changes after simultaneously GARP and EARP depletion.

### GARP and EARP specific effects on lysosomal homeostasis

To further relate our findings directly to GARP or EARP function, we used the VPS54-KD to selectively disrupt GARP complex function. We observed a de-enrichment of proteins with primarily lysosomal-related GO terms when comparing VPS54-KD lysosomes to control lysosomes (Fig. 5c, d). A phenotype also observed in the VPS53 knockdown (vacuolar membrane, lysosomal membrane, lytic vacuole membrane) (Fig. 5c, d). This included the de-enrichment of NAGAB, two pore channel protein 1 (TPC1), lysosomal cobalamin transporter (ABCD4), Sortilin-related receptor (SORL), amino acid transporter (LAT1), lysosomal serine protease TPP1, PPT1, and LIPL. Additionally, Golgi-related proteins were also de-enriched in VPS54-KD lysosomes as found by GO term analysis with the terms trans-Golgi network and Golgi apparatus (Fig. 5d). To exclude a co-enrichment of Golgi fractions in our lysosomal pull-downs, we checked for the Golgi marker protein (GM130) in Lyso-IPs of VPS54-KD, VPS53-KD and control cells via western blotting (Fig. S3b, c). Good enrichment of the bait protein LAMP1 and GFP, but no co-enrichment of GM130 excluded this possibility. Together, these results demonstrate diminished lysosomal delivery of luminal proteins in the absence of GARP, which are presumably retained in the Golgi apparatus instead.

EARP specific knockdown of VPS50 resulted in an increase of extracellular and endosomal proteins in the lysosome, such as collagen chain CO3A1, sorting nexin SNX8, LIPL, metalloprotease STABP, and dermcidin (DCD). This was further underlined by the GO term analysis, which identified early endosomes, focal adhesion, collagen-containing extracellular matrix, and cell-substrate junctions. This effect was already observed in VPS53KD lysosomes and indicates that this finding can be directly related to EARP dysfunction. De-enriched lysosomal proteins in the EARP complex mutant were cytosolic and ribosomal proteins such as pyruvate kinase KPYR, Glucocorticoid receptor (GCR), protein disulfide isomerase CREL1 and ribosomal RNA processing protein RRP1 (Fig. 5e, f). The lower gene counts associated with the GO terms following VPS50 depletion compared to a VPS53 depletion likely reflect the previously observed partial functional compensation by the intact GARP complex. In contrast, depletion of the shared subunit VPS53 disrupts both complexes, eliminating this compensatory capacity and resulting in a broader proteomic phenotype (Gershlick et al., 2019; Schindler et al., 2015). Alternatively, the more pronounced enrichment may indicate that GARP itself contributes directly to these pathways.

### GARP and EARP Loss Induces Distinct Cargo Misrouting Patterns

To further characterize and confirm the mass spectrometry-based findings, we chose three clusters from the heat map in Fig. 4d (green boxes) and selected one exemplary missorted cargo from each cluster (Fig. 6a; PPT1, NAGAB and LIPL). We first confirmed their lysosomal enrichment and de-enrichment by western blotting. PPT1 demonstrated the same tendency as observed in the proteomics data with a decrease in lysosomes in VPS52KD, VPS53KD and unchanged levels in VPS54-KD compared to control Lyso-IPs (Fig. 5b, S5a, 6b). This is also true for LIPL which remained unchanged in the lysosomes after VPS50-KD, but increased in VPS52-KD and VPS53-KO lysosomes as observed by mass spectrometry (Fig. 5b, S5a, 6b). These findings were further supported by confocal microscopy of VPS50-KD, VPS53-KD and VPS54-KD knockdown cells, combined with immunostaining for PPT1 and NAGAB (Fig. 6c). Here, colocalization with LAMP1 as lysosomal marker was decreased after GARP and simultaneous GARP and EARP depletion, but not by depleting the EARP-specific subunit VPS50. Colocalization of PPT1 with the Golgi signal of TGN46 remained unaltered. Colocalization of a NAGAB-specific antibody with LAMP1 was lost after VPS53-KD and VPS54-KD, but not after VPS50-KD as shown by microscopy and mass spectrometry (Fig. 6c). Instead, NAGAB colocalizing with the Golgi marker TGN46 was significantly increased in a VPS53 deletion (Fig. S5b and c). For LIPL, we found increased colocalization with LAMP1 in VPS50-KD and VPS53-KD cells, but not in VPS54-KD cells (Fig. 6c). This further confirmed the mis-sorting of extracellular proteins to the lysosomes without functional EARP complex. At the same time, colocalization of LIPL with the Golgi apparatus (TGN46) was slightly increased in the VPS53-KD lysosomes (Fig. S5b and c). Taken together, these confocal microscopy and immunoblotting data recapitulated our findings from mass spectrometry-based proteomics of lysosomes combined with acute downregulation of trafficking complexes.

**Fig. 6:**
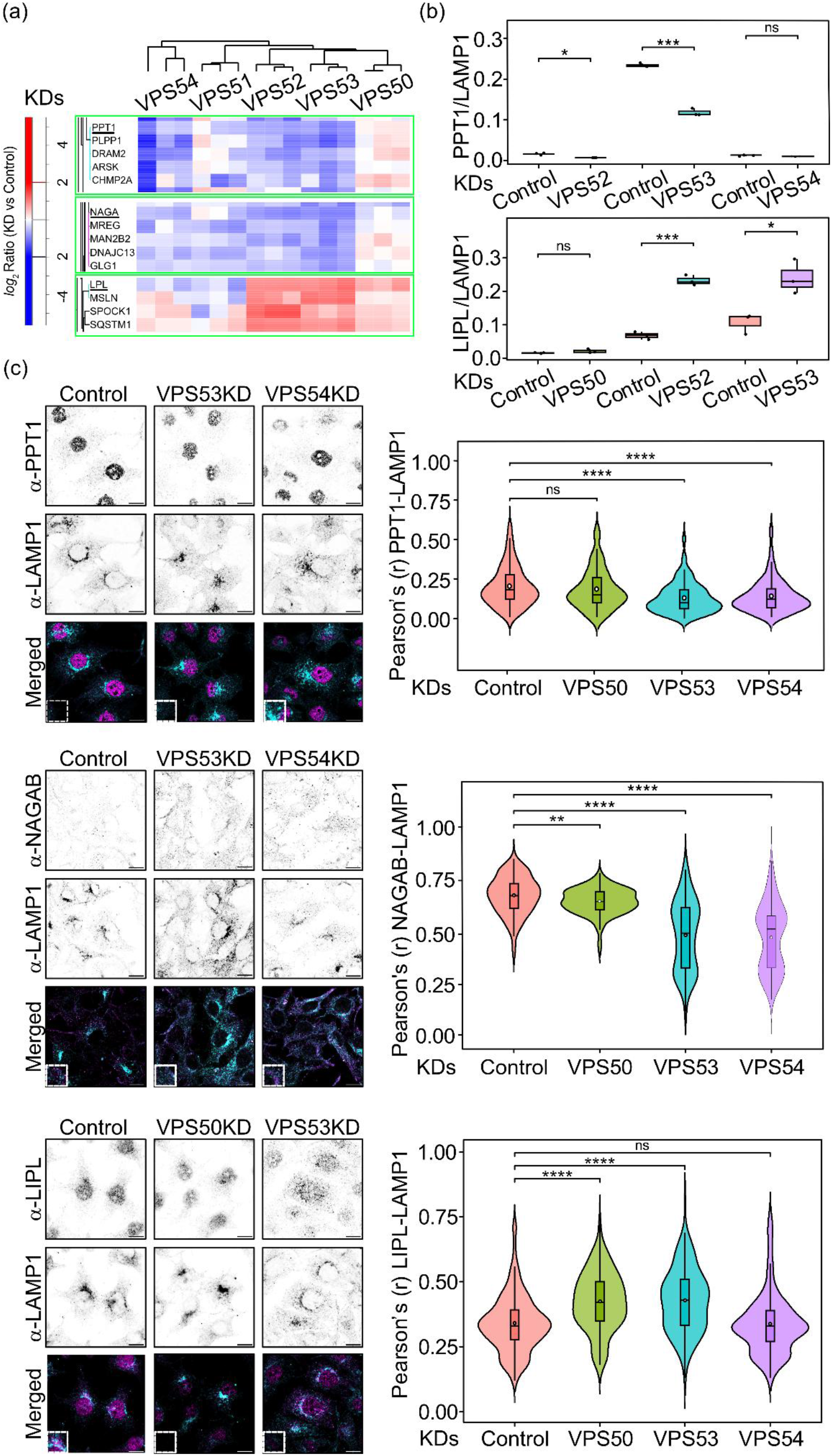
Validation of selected protein effects upon disruptive retrograde and recycling pathways. (a) Clustering analysis using a heatmap showing fold changes of selected proteins in Lyso-IP from GARP/EARP knockdowns relative to control Lyso-IPs. Proteins de-enriched in knockdown lysosomes (PPT1 and NAGAB) are shown in blue, while enriched proteins (LIPL) are shown in red, with gradient colors reflecting the magnitude of fold change. (b) Western blot quantification of PPT1 and LIPL (normalized to LAMP1) in control and knockdown lysosomes, represented as box plots with statistical significance indicated by stars (*p < 0.05, **p < 0.01, ***p < 0.001) (n=3), demonstrating impaired lysosomal enzyme targeting and enhanced cargo retention. (c) Immunostaining of PPT1, NAGAB, and LIPL with LAMP1 in control and GARP/EARP knockdown cells (*Scale bar: 10 µm*), with colocalization quantified and displayed as violin plots (right). Statistical significance is indicated by stars (*p < 0.05, **p < 0.01, ***p < 0.001, ****p < 0.0001). (n = 3, with n ≥ 104 cells quantified per replicate).

## Discussion

Multi-subunit tethering complexes are important to maintain endo-lysosomal sorting in cells, with the GARP and EARP complexes having important roles in endosomal recycling (Pérez-Victoria et al., 2008; Schindler et al., 2015). While these complexes are extensively studied regarding certain model cargoes, a general understanding of the broad spectrum of cargoes is largely lacking. This is especially important since the observed phenotypes in KOs or natural occurring mutants can generally not be attributed to a single cause. Thus, systematic analyses of the phenotypes are highly relevant to understand the complex physiology in disease-causing mutations.

Here, we established a new protocol to purify lysosomes from LAMP1-GFP expressing HeLa cells. Our in-depth analysis of the lysosomal proteome and cross-validation with similar datasets (Akter et al., 2023; Hein et al., 2025) revealed the high quality of our lysosomal purifications. We provide a comprehensive list of lysosomal proteins in the HeLa cell background used here and carefully evaluate the selected cutoffs for defining lysosomal proteins. Furthermore, we used siRNA KDs of the two mannose-6-phosphate receptors to define the dynamic range of lysosomal proteome variations upon knockdown of genes. Although, we detected relatively small changes, we consider them highly relevant as they directly reflect early occurring changes, giving the cells little time to adapt. While KD efficiencies are around 70-80%, the remaining protein amounts are probably sufficient to maintain some basic protein levels and therefore some remaining functionality. A similar approach in yeast, where we used the auxin-induced degron system, allowed us to identify early-onset changes in the abundance of aminophospholipid flippases in GARP knockdowns (Eising et al., 2019).

After benchmarking our lysosomal proteome and confirming the validity of siRNA-mediated knockdowns followed by lysosomal proteomics, we characterized the lysosomal proteome of knockdowns of all GARP and EARP subunits. This is highly relevant as mutations in GARP and EARP subunits are directly linked to the development of several neurodegenerative disorders (Feinstein et al., 2014; Gershlick et al., 2019; Schneeberger et al., 2021). As expected, we observed the accumulation of various proteins in the purified lysosome fraction. This fraction correlates well with a previous study investigating the Golgi proteome in the wobbler mouse (Petit et al., 2020) harboring a mutation in the Vps54 subunit. Interestingly, the GARP-specific effects in the VPS54KD were more severe compared to the EARP-specific VPS50KD. Why are many proteins accumulating at lysosomes in a GARP KD? We speculate the retrograde protein carriers that are not tethered at the Golgi ultimately end up in the lysosome. This can be explained by a model where retrograde transport carriers remain tubular, while tethering and fusion at the Golgi occur simultaneously with tubule fission at endosomes. If the Golgi tethering machinery is affected, the cargoes remain in endosomes and are ultimately delivered to lysosomes. Other proteins are depleted in the lysosomes of GARP mutants which might be explained by the loss of sorting receptors, necessary to sort these proteins into the endo-lysosomal pathway (Bonifacino and Hierro, 2011; Ghosh et al., 2026; Simonetti et al., 2017). The correlation between the different KDs of the GARP and EARP subunits is relatively low. Amongst the different KDs, VPS51KD shows the mildest effects which has also been observed in other studies in both yeast and mammalian cells (Conibear et al., 2003; Luo et al., 2011; Siniossoglou and Pelham, 2002).

Our analysis revealed that multiple enzymes involved in lipid metabolism are mis-sorted in knockdowns of the GARP and EARP complex. While some of the changes are relatively mild, we could confirm several of the hits independently using confocal microscopy and western blot assays. Amongst the depleted protein hits is also the cholesterol and sphingolipid transporting protein NPC2 which is in line with literature showing accumulation of sterols and sphingolipids in GARP mutants (Bonifacino and Hierro, 2011; Eising et al., 2019; Wei et al., 2017). However, the effects of mutations in GARP and EARP are difficult to narrow down to a single phenotype. Our data support a model in which diseases associated with the depletion of GARP or EARP subunits are more systemic. Multiple proteins involved in lipid metabolism are mis-localized in cells, suggesting that lipid metabolism in general is altered. Going forward, this highlights the need for systematic methods to study the effects of mutations in the endo-lysosomal system, which typically cannot be attributed to a single phenotype.

## Materials and methods

HeLa cells were cultured in Dulbecco’s modified Eagle’s medium (DMEM) supplemented with 10% fetal bovine serum (FBS; Pan Biotech) at 37 °C in a humidified atmosphere containing 5% CO_2_, unless otherwise specified.

### Immunoprecipitation of intact lysosomes (Lyso-IP)

HeLa wild-type (WT) and LAMP1-GFP expressing cells were cultured in 15-cm dishes (∼80% confluency). Cells were scraped in ice-cold PBS, pelleted (500 g, 5 min, 4 °C), and washed twice with lysis buffer 1 (210 mM mannitol, 70 mM sucrose, 10 mM HEPES, pH 7.5) by centrifugation (1000 g, 5 min, 4 °C) without resuspension. The pellet was gently resuspended in 800 µL lysis buffer 2 (lysis buffer 1 supplemented with 0.5 mM DTT, protease inhibitor cocktail, and DNase I) and homogenized on ice using a Balch homogenizer (25 strokes). Lysates were centrifuged twice (1500 g, 4 °C) to remove nuclear debris (10 min) and light membranes (15 min), and the post-nuclear supernatant (PNS) was collected.

Protein concentrations were determined using a Bradford assay. For Lyso-IP, samples were adjusted to 500 µg protein, and 5 mg/ml fatty acid-free BSA was added. Samples were incubated with 25 µl equilibrated anti-GFP microbeads (µMACS GFP Isolation Kit, Miltenyi Biotec) for 1 h at 4 °C with gentle rotation. Bead-lysate mixtures were applied to pre-equilibrated µColumns (Miltenyi Biotec) on a µMACS Separator (Miltenyi Biotec), washed once with lysis buffer 3 (lysis buffer 2 adjusted to 5 mg/ml BSA) and three times with lysis buffer 1. The beads were eluted in 600 µL lysis buffer 1 and pelleted (21000 g, 20 min, 4 °C). GFP-bound organelles were eluted under denaturing conditions (0.5% SDS in lysis buffer 1, 10 min, RT), and supernatants were collected as immunoprecipitated (IP) samples. IP fractions (20 µL) and 10 µg of PNS were lysed, digested, and processed using the PreOmics iST or S-Trap^TM^ micro sample prep kit and analyzed by LC-MS/MS.

### Generation of Knockdown mutants

HeLa wild-type (WT) and LAMP1-GFP expressing cells were transfected with ON-TARGETplus siRNAs (Dharmacon™, Horizon Discovery) targeting VPS50, VPS51, VPS52, VPS53, VPS54, cation-dependent M6PR (M6PR), or cation-independent M6PR (IGF2R). Approximately 40% confluent 10-cm dishes of HeLa-WT and HeLa-LAMP1-GFP cells were transfected with either non-targeting control or target-specific siRNAs using Fugene® SI according to the manufacturer’s protocol. Briefly, siRNA-Fugene complexes were prepared in serum-free OptiMEM™ (Gibco™) and added dropwise to the culture medium (final siRNA concentration: 150 pmol per dish). Cells were incubated for 72 h at 37 °C with 5% CO₂. After 68 - 72 h, cells were expanded into triplicate 15-cm dishes and subjected to a second round of transfection under identical conditions (225 pmol siRNA per dish). Following an additional 72-h incubation, knockdown efficiency was validated via western blot, and cells were harvested for LysoIP and downstream proteomic analysis.

### Western blot

For immunoblotting of LysoIP, HeLa wild-type (WT) and LAMP1-GFP cells, with or without GARP/EARP knockdowns, were cultured to ∼80% confluency in 10-cm dishes. Lysosome immunoprecipitation was performed as described above, and the bead-bound fractions were eluted with Laemmli buffer containing DTT at 65 °C for 10 min. After centrifugation, the supernatants were analyzed by SDS-PAGE and immunoblotting. GFP-tagged proteins were detected using a 1:1000 dilution of mouse anti-GFP antibody (11814460001, Sigma-Aldrich). Additional antibodies included 1:1000 dilution of rabbit anti-LAMP1 (9091, Cell Signaling Technology), anti-Rab7 (9367, Cell Signaling Technology), and anti-Rab5 (3547, Cell Signaling Technology), mouse anti-tubulin (3873, Cell Signaling Technology), mouse anti-mitoP60 (Mab1273, Millipore), and mouse anti-calnexin (23954, Santa Cruz Biotechnology). Secondary antibodies conjugated with DyLight^TM^ 800, goat anti-rabbit (SA535571, Invitrogen) and goat anti-mouse (SA535521, Invitrogen), in 1:20000 dilution were detected using ChemiDoc MP (Bio-Rad).

To validate knockdown efficiencies, HeLa WT and LAMP1-GFP cells were transfected twice with non-targeting control or target-specific siRNAs as described above. Cells were lysed in RIPA buffer (150 mM NaCl, 25 mM Tris-HCl pH 7.6, 1% NP-40, 1% sodium deoxycholate, 0.1% SDS) supplemented with protease inhibitor cocktail (cOmplete ULTRA Tablets, Roche) and DNase I (PanReac AppliChem). Lysates were centrifuged (600 g, 10 min, 4 °C), and the PNS was mixed with Laemmli buffer containing DTT and boiled at 95 °C for 10 min. Western blots were probed with mouse anti-VPS50 (1:500, H00055610-M01, Abnova), 1:1000 rabbit anti-VPS51 (HPA039650, Atlas Antibodies) and anti-VPS53 (HPA024446, Atlas Antibodies), rabbit anti-VPS54 (1:1000, STJ115181, St. John’s Laboratory), and rabbit anti-IGF2R (1:1000, 14364, Cell Signaling Technology). HRP-conjugated 1:10000 goat anti-mouse (31430, Thermo Fischer Scientific) and mouse anti-rabbit (2357, Santa Cruz Biotechnology) were used for chemiluminescent detection.

Western blot signals were quantified using *Image Lab* software (ChemiDoc MP, Bio-Rad). Bands of interest were manually defined, and adjusted volume intensity (Int) values were measured. Background subtraction was applied for each lane, and signal intensities were normalized to the corresponding loading controls. Normalized values were exported and visualized as box plots to compare protein expression levels across knockdown conditions.

### Mass Spectrometric Data Acquisition and Analysis

Immunoprecipitated proteins were processed using the PreOmics iST or S-Trap^TM^ micro sample preparation kit according to the manufacturer’s instructions for downstream LC-MS/MS analysis. Data-independent acquisition (DIA) was performed for label-free quantification (LFQ).

Dried peptides were resuspended in LC-LOAD buffer, and 1–2 µl (≈100 ng protein) was injected for reversed-phase chromatography on a Thermo Ultimate 3000 RSLCnano system coupled to a TimsTOF HT mass spectrometer (Bruker Corporation, Bremen, Germany) via a CaptiveSpray Ion source. Peptides were separated on an Aurora Gen3 C18 column (25 cm × 75 µm, 1.6 µm particle size) with an CSI emitter (IonOpticks, Australia) at 40 °C. Elution of peptides from the column was realized via a linear gradient of acetonitrile from 2-35% (in water with 0.1% formic acid) for 44 min at a constant flow rate of 300 nL/min following a 7 min increase to 50%, and finally, 4 min to reach 85% and electrosprayed at 1.5 kV with a dry gas flow of 3 l/min. The TimsTOF HT was operated in positive Ion mode (scan range: 100–1700 m/z) with an Ion mobility ramp of 0.7–1.5 Vs/cm² over 100 ms. The accumulation time was also set to 100 ms. The scan mode was set to DIA-PASEF. 10 MS1 ramps with a window of 26 Da were selected within the mass range 284 to 1385 and the mobility range from 0.7 to 1. 42 [1/k0], resulting in 44 steps per cycle with a mass overlap of 1 Da.

Raw data were processed with MaxQuant (v2.6.7.0; www.maxquant.org) (Cox et al., 2011; Cox and Mann, 2008). Quantification was performed using ID-directed LFQ. Protein group outputs from MaxQuant were processed in Perseus (v2.0.11; www.maxquant.org/perseus) (Tyanova et al., 2016). Contaminants and reverse hits were removed, LFQ intensities were *log*_2_-transformed, and missing values were imputed from a normal distribution. Differential enrichment between conditions was assessed by two-sided Student’s t-test, visualized as volcano plots showing *log*_2_ Ratio versus -*log*_10_ P-value.

### Enrichment and Clustering Analysis

Significantly enriched proteins were subjected to Gene Ontology (GO) enrichment analysis in R (v4.4.3) using clusterProfiler (Yu et al., 2012). Gene names were mapped to Entrez IDs via the bitr() function referencing org.Hs.eg.db. GO enrichment was performed with enrichGO() using the “cellular component” ontology and Benjamini-Hochberg (BH) correction. The top enriched GO terms were visualized as bar plots generated with ggplot2 (Wickham, 2016).

Hierarchical clustering of *log*_2_-transformed LFQ intensities was performed in Perseus using Euclidean distance and average linkage. Protein and sample values were z-scored, and missing data were imputed (width = 0.3, downshift = 1.8). Clustering results were visualized as heatmaps to identify proteins with similar abundance profiles across knockdown conditions.

Additionally, soft clustering was conducted using the Mfuzz package (Futschik and Carlisle, 2005) in R to detect co-regulated protein modules. Data were filtered, *log*2-transformed, and standardized. Optimal fuzzification (m) and cluster number (c) were estimated with mestimate() and cselection(), and clusters were visualized using mfuzz.plot() to reveal gradual or overlapping abundance trends.

### Immunofluorescence and Confocal Microscopy

Candidate proteins (PPT1, NAGAB, and LIPL) identified from the proteomic analysis were immunostained together with the lysosomal marker, LAMP1, and the Golgi marker, TGN46. Immunostaining was performed under control and gene knockdown conditions. HeLa-LAMP1-GFP cells were transfected with control or target-specific siRNAs as described before. After 48 h, approximately 5 × 10⁴ cells were seeded onto sterile 12-mm glass coverslips in 24-well plates, followed by a second siRNA transfection and 24 h incubation at 37 °C with 5% CO₂.

Cells were fixed with 4% paraformaldehyde (PFA) in PBS for 10 min at room temperature (RT), washed with PBS, and permeabilized with 0.3% Triton X-100 and 0.3% SDS in PBS for 15 min at RT. After washing, samples were blocked with 2% BSA in PBS for 1 h at RT. Coverslips were incubated overnight at 4 °C with primary antibodies diluted in 1% BSA and 0.3% Triton X-100 in PBS: rabbit anti-LIPL (1:100, ATA-HPA048749, Biozol), rabbit anti-PPT1 (1:400, HPA021546, Sigma-Aldrich), rabbit anti-NAGAB (1:50, 15096-1-AP, Proteintech), mouse anti-LAMP1 (1:400, sc-20011, Santa Cruz), and sheep anti-TGN46 (1:400, AHP500GT, Bio-Rad). After washing, cells were incubated for 1 h at RT with 1:250 diluted fluorophore-conjugated secondary antibodies: anti-rabbit Cy^TM^3 and anti-mouse Cy^TM^5 (Jackson ImmunoResearch Laboratories), and anti-sheep/goat DyLight 488 (Bio-Rad).

Following final washes with PBS and distilled water, coverslips were mounted using ProLong™ Gold Antifade Mountant (Thermo Fisher Scientific) and dried overnight at RT. Confocal images were acquired using a Zeiss LSM 880 microscope with a 63x oil objective, equipped with an Airyscan detector under ambient temperature conditions.

### Image Analysis and Quantification

Confocal images were analyzed using *ImageJ/Fiji* 1.54g (Schindelin et al., 2012) to quantify the colocalization of LIPL, NAGAB, and PPT1 with the lysosomal marker LAMP1 and the Golgi marker TGN46. Regions of interest (ROIs) were manually defined for individual cells, and colocalization was evaluated using the Coloc2 plugin (Bolte and Cordelières, 2006). Pearson’s correlation coefficient (PCC) was calculated to assess the degree of spatial overlap between target proteins and organelle markers. Acquired raw images were processed in Fiji, and quantitative data were visualized as violin plots in R (v4.4.3).

### Data availability

All proteomics data are available in Supplemental Table 1. The mass spectrometry proteomics data have been deposited to the ProteomeXchange Consortium via the PRIDE partner repository with the dataset identifier PXD084738 (Deutsch et al., 2026; Perez-Riverol et al., 2025).

## Acknowledgements

We thank members of the Fröhlich lab for careful reading and commenting on the manuscript. Florian Fröhlich is a member of the Heisenberg program of the DFG (491484150). Work in the Fröhlich lab is supported by the SFB1557 project Z1 (516911785), P6 (467522186), the DFG (534958928), (282452898), (503478512) and the Boehringer Ingelheim Foundation (RISE UP! program).

## Notes

### Competing Interest Statement

The authors have declared no competing interest.

### Summary of Updates

The previous version of the manuscript was uploaded without supplementary material via review commons. Now, A version without the Review Commons page but with the supplementary material included was uploaded.

## References

1. Abu-Remaileh M, Wyant GA, Kim C, Laqtom NN, Abbasi M, Chan SH, Freinkman E, Sabatini DM. 2017. Lysosomal metabolomics reveals V-ATPase- and mTOR-dependent regulation of amino acid efflux from lysosomes. Science 358:807–813. DOI: 10.1126/science.aan6298

2. Adeosun OA, Schröer C, Südhoff E, Bergenthal F, Sommer K, Döffinger E, Fiedler B, Hilderink A, Lehmann A-K, Pohle LS, Korneev SM, Fröhlich F, Maeda K, Kurre R, Holtmannspötter M, Esch BM, Holthuis JCM. 2026. VPS13C/PARK23 initiates lipid transfer and membrane remodeling for efficient lysosomal repair. Nature Communications 17:5789. DOI: 10.1038/s41467-026-75145-y

3. Akter F, Bonini S, Ponnaiyan S, Kögler-Mohrbacher B, Bleibaum F, Damme M, Renard BY, Winter D. 2023. Multi–Cell Line Analysis of Lysosomal Proteomes Reveals Unique Features and Novel Lysosomal Proteins. Molecular C Cellular Proteomics 22:100509. DOI: 10.1016/j.mcpro.2023.100509

4. Aragon-Ramirez WS, Khakurel A, Pokrovskaya I, Sumya FT, Lupashin VV. 2026. Golgi CATCHR complexes function as organizing hubs for vesicle tethering and fusion. DOI: 10.64898/2026.06.16.732723

5. Arighi CN, Hartnell LM, Aguilar RC, Haft CR, Bonifacino JS. 2004. Role of the mammalian retromer in sorting of the cation-independent mannose 6-phosphate receptor. The Journal of Cell Biology 165:123–133. DOI: 10.1083/jcb.200312055

6. Beilina A, Bonet-Ponce L, Kumaran R, Kordich JJ, Ishida M, Mamais A, Kaganovich A, Saez-Atienzar S, Gershlick DC, Roosen DA, Pellegrini L, Malkov V, Fell MJ, Harvey K, Bonifacino JS, Moore DJ, Cookson MR. 2020. The Parkinson’s Disease Protein LRRK2 Interacts with the GARP Complex to Promote Retrograde Transport to the trans-Golgi Network. Cell Reports 31:107614. DOI: 10.1016/j.celrep.2020.107614

7. Bolte S, Cordelières FP. 2006. A guided tour into subcellular colocalization analysis in light microscopy. Journal of Microscopy 224:213–232. DOI: 10.1111/j.1365-2818.2006.01706.x

8. Bonifacino JS, Hierro A. 2011. Transport according to GARP: receiving retrograde cargo at the trans-Golgi network. Trends in Cell Biology 21:159–167. DOI: 10.1016/j.tcb.2010.11.003

9. Bonifacino JS, Rojas R. 2006. Retrograde transport from endosomes to the trans-Golgi network. Nature Reviews Molecular Cell Biology 7:568–579. DOI: 10.1038/nrm1985

10. Carlton J, Bujny M, Rutherford A, Cullen P. 2005. Sorting Nexins – Unifying Trends and New Perspectives. Traffic 6:75–82. DOI: 10.1111/j.1600-0854.2005.00260.x

11. Conibear E, Cleck JN, Stevens TH. 2003. Vps51p Mediates the Association of the GARP (Vps52/53/54) Complex with the Late Golgi t-SNARE Tlg1p. Molecular Biology of the Cell 14:1610–1623. DOI: 10.1091/mbc.e02-10-0654

12. Cox J, Mann M. 2008. MaxQuant enables high peptide identification rates, individualized p.p.b.-range mass accuracies and proteome-wide protein quantification. Nature Biotechnology 26:1367–1372. DOI: 10.1038/nbt.1511

13. Cox J, Neuhauser N, Michalski A, Scheltema RA, Olsen JV, Mann M. 2011. Andromeda: A Peptide Search Engine Integrated into the MaxQuant Environment. Journal of Proteome Research 10:1794–1805. DOI: 10.1021/pr101065j

14. Cullen PJ. 2008. Endosomal sorting and signalling: an emerging role for sorting nexins. Nature Reviews Molecular Cell Biology 9:574–582. DOI: 10.1038/nrm2427

15. Deutsch EW, Bandeira N, Perez-Riverol Y, Sharma V, Carver JJ, Mendoza L, Kundu DJ, Bandla C, Kamatchinathan S, Hewapathirana S, Sun Z, Kawano S, Okuda S, Connolly B, MacLean B, MacCoss MJ, Chen T, Zhu Y, Ishihama Y, Vizcaíno JA. 2026. The ProteomeXchange consortium in 2026: making proteomics data FAIR. Nucleic Acids Research 54:D459–D469. DOI: 10.1093/nar/gkaf1146

16. Duchen LW, Strich SJ. 1968. An hereditary motor neurone disease with progressive denervation of muscle in the mouse: the mutant “wobbler”. Journal of Neurology, Neurosurgery C Psychiatry 31:535–542. DOI: 10.1136/jnnp.31.6.535

17. Eising S, Esch B, Wälte M, Vargas Duarte P, Walter S, Ungermann C, Bohnert M, Fröhlich F. 2022. A lysosomal biogenesis map reveals the cargo spectrum of yeast vacuolar protein targeting pathways. Journal of Cell Biology 221:e202107148. DOI: 10.1083/jcb.202107148

18. Eising S, Thiele L, Fröhlich F. 2019. A systematic approach to identify recycling endocytic cargo depending on the GARP complex. eLife 8:e42837. DOI: 10.7554/eLife.42837

19. Feinstein M, Flusser H, Lerman-Sagie T, Ben-Zeev B, Lev D, Agamy O, Cohen I, Kadir R, Sivan S, Leshinsky-Silver E, Markus B, Birk OS. 2014. *VPS53* mutations cause progressive cerebello-cerebral atrophy type 2 (PCCA2). Journal of Medical Genetics 51:303–308. DOI: 10.1136/jmedgenet-2013-101823

20. Fröhlich F, Petit C, Kory N, Christiano R, Hannibal-Bach H-K, Graham M, Liu X, Ejsing CS, Farese RV, Walther TC. 2015. The GARP complex is required for cellular sphingolipid homeostasis. eLife 4:e08712. DOI: 10.7554/eLife.08712

21. Futschik ME, Carlisle B. 2005. Noise-robust soft clustering of gene expression time-course data. Journal of Bioinformatics and Computational Biology 03:965–988. DOI: 10.1142/S0219720005001375

22. Gauthier C, El Cheikh K, Basile I, Daurat M, Morère E, Garcia M, Maynadier M, Morère A, Gary-Bobo M. 2024. Cation-independent mannose 6-phosphate receptor: From roles and functions to targeted therapies. Journal of Controlled Release 365:759–772. DOI: 10.1016/j.jconrel.2023.12.014

23. Gershlick DC, Ishida M, Jones JR, Bellomo A, Bonifacino JS, Everman DB. 2019. A neurodevelopmental disorder caused by mutations in the VPS51 subunit of the GARP and EARP complexes. Human Molecular Genetics 28:1548–1560. DOI: 10.1093/hmg/ddy423

24. Ghosh S, Singh J, Damseh NS, Severino M, De Pace R, Golding AE, Jarnik M, Thakran P, Faivre L, Heitz J, Denommé-Pichon A-S, Vitobello A, AlAbdi L, Abdulwahab F, Sumayli S, Alqahtani M, Cheema HA, Javed I, Kim J, Lochmüller H, Mor-Shaked H, Neil JE, Mochida GH, Zifarelli G, Bauer P, Barkhordari E, Ghayoor Karimiani E, Houlden H, Abu-Libdeh B, Edvardson S, Elpeleg O, Maroofian R, Patten SA, Bonifacino JS. 2026. *EIPR1* variants cause a neurodevelopmental disorder with endolysosomal and dense core vesicle defects. Brain 149:1568–1585. DOI: 10.1093/brain/awaf371

25. Hein MY, Peng D, Todorova V, McCarthy F, Kim K, Liu C, Savy L, Januel C, Baltazar-Nunez R, Sekhar M, Vaid S, Bax S, Vangipuram M, Burgess J, Njoya L, Wang E, Ivanov IE, Byrum JR, Pradeep S, Gonzalez CG, Aniseia Y, Creery JS, McMorrow AH, Sunshine S, Yeung-Levy S, DeFelice BC, Mehta SB, Itzhak DN, Elias JE, Leonetti MD. 2025. Global organelle profiling reveals subcellular localization and remodeling at proteome scale. Cell 188:1137–1155.e20. DOI: 10.1016/j.cell.2024.11.028

26. Hirata T, Fujita M, Nakamura S, Gotoh K, Motooka D, Murakami Y, Maeda Y, Kinoshita T. 2015. Post-Golgi anterograde transport requires GARP-dependent endosome-to-TGN retrograde transport. Molecular Biology of the Cell 26:3071–3084. DOI: 10.1091/mbc.E14-11-1568

27. Hong W, Lev S. 2014. Tethering the assembly of SNARE complexes. Trends in Cell Biology 24:35–43. DOI: 10.1016/j.tcb.2013.09.006

28. Khakurel A, Kudlyk T, Bonifacino JS, Lupashin VV. 2021. The Golgi-associated retrograde protein (GARP) complex plays an essential role in the maintenance of the Golgi glycosylation machinery. Molecular Biology of the Cell 32:1594–1610. DOI: 10.1091/mbc.E21-04-0169

29. Khakurel A, Kudlyk T, Pokrovskaya I, D’Souza Z, Lupashin VV. 2022. GARP dysfunction results in COPI displacement, depletion of Golgi v-SNAREs and calcium homeostasis proteins. Frontiers in Cell and Developmental Biology 10:1066504. DOI: 10.3389/fcell.2022.1066504

30. Khakurel A, Pokrovskaya I, Aragon-Ramirez WS, Lupashin VV. 2025. Acute GARP Depletion Disrupts Vesicle Transport, Leading to Severe Defects in Sorting, Secretion and *O* -Glycosylation. Traffic 26:e70003. DOI: 10.1111/tra.70003

31. Kumar L, Futschik ME. 2007. Mfuzz: A software package for soft clustering of microarray data. Bioinformation 2:5–7. DOI: 10.6026/97320630002005

32. Kümmel D, Ungermann C. 2014. Principles of membrane tethering and fusion in endosome and lysosome biogenesis. Current Opinion in Cell Biology 29:61–66. DOI: 10.1016/j.ceb.2014.04.007

33. Liewen H, Meinhold-Heerlein I, Oliveira V, Schwarzenbacher R, Luo G, Wadle A, Jung M, Pfreundschuh M, Stenner-Liewen F. 2005. Characterization of the human GARP (Golgi associated retrograde protein) complex. Experimental Cell Research 306:24–34. DOI: 10.1016/j.yexcr.2005.01.022

34. Luo L, Hannemann M, Koenig S, Hegermann J, Ailion M, Cho M-K, Sasidharan N, Zweckstetter M, Rensing SA, Eimer S. 2011. The *Caenorhabditis elegans* GARP complex contains the conserved Vps51 subunit and is required to maintain lysosomal morphology. Molecular Biology of the Cell 22:2564–2578. DOI: 10.1091/mbc.e10-06-0493

35. Malerød L, Pedersen NM, Sem Wegner CE, Lobert VH, Leithe E, Brech A, Rivedal E, Liestøl K, Stenmark H. 2011. Cargo-Dependent Degradation of ESCRT-I as a Feedback Mechanism to Modulate Endosomal Sorting. Traffic 12:1211–1226. DOI: 10.1111/j.1600-0854.2011.01220.x

36. Perez-Riverol Y, Bandla C, Kundu DJ, Kamatchinathan S, Bai J, Hewapathirana S, John NS, Prakash A, Walzer M, Wang S, Vizcaíno JA. 2025. The PRIDE database at 20 years: 2025 update. Nucleic Acids Research 53:D543–D553. DOI: 10.1093/nar/gkae1011

37. Pérez-Victoria FJ, Abascal-Palacios G, Tascón I, Kajava A, Magadán JG, Pioro EP, Bonifacino JS, Hierro A. 2010. Structural basis for the wobbler mouse neurodegenerative disorder caused by mutation in the Vps54 subunit of the GARP complex. Proceedings of the National Academy of Sciences 107:12860–12865. DOI: 10.1073/pnas.1004756107

38. Pérez-Victoria FJ, Mardones GA, Bonifacino JS. 2008. Requirement of the Human GARP Complex for Mannose 6-phosphate-receptor-dependent Sorting of Cathepsin D to Lysosomes. Molecular Biology of the Cell 19:2350–2362. DOI: 10.1091/mbc.e07-11-1189

39. Petit CS, Lee JJ, Boland S, Swarup S, Christiano R, Lai ZW, Mejhert N, Elliott SD, McFall D, Haque S, Huang EJ, Bronson RT, Harper JW, Farese RV, Walther TC. 2020. Inhibition of sphingolipid synthesis improves outcomes and survival in GARP mutant *wobbler* mice, a model of motor neuron degeneration. Proceedings of the National Academy of Sciences 117:10565–10574. DOI: 10.1073/pnas.1913956117

40. Raiborg C, Stenmark H. 2009. The ESCRT machinery in endosomal sorting of ubiquitylated membrane proteins. Nature 458:445–452. DOI: 10.1038/nature07961

41. Saarela D, Lis P, Gomes S, Nirujogi RS, Dong W, Rawat E, Glendinning S, Zeneviciute K, Bagnoli E, Fasimoye R, Lin C, Nyame K, Boros FA, Zunke F, Lamoliatte F, Elshani S, Jaconelli M, Jans JJM, Huisman MA, Posern C, Westermann LM, Schulz A, Van Hasselt PM, Alessi DR, Abu-Remaileh M, Sammler EM. 2024. Tagless LysoIP method for molecular profiling of lysosomal content in clinical samples. DOI: 10.1101/2024.05.17.594681

42. Schindelin J, Arganda-Carreras I, Frise E, Kaynig V, Longair M, Pietzsch T, Preibisch S, Rueden C, Saalfeld S, Schmid B, Tinevez J-Y, White DJ, Hartenstein V, Eliceiri K, Tomancak P, Cardona A. 2012. Fiji: an open-source platform for biological-image analysis. Nature Methods 9:676– 682. DOI: 10.1038/nmeth.2019

43. Schindler C, Chen Y, Pu J, Guo X, Bonifacino JS. 2015. EARP is a multisubunit tethering complex involved in endocytic recycling. Nature Cell Biology 17:639–650. DOI: 10.1038/ncb3129

44. Schmitt-John T, Drepper C, Mußmann A, Hahn P, Kuhlmann M, Thiel C, Hafner M, Lengeling A, Heimann P, Jones JM, Meisler MH, Jockusch H. 2005. Mutation of Vps54 causes motor neuron disease and defective spermiogenesis in the wobbler mouse. Nature Genetics 37:1213–1215. DOI: 10.1038/ng1661

45. Schneeberger PE, Nampoothiri S, Holling T, Yesodharan D, Alawi M, Knisely AS, Müller T, Plecko B, Janecke AR, Kutsche K. 2021. Biallelic variants in *VPS50* cause a neurodevelopmental disorder with neonatal cholestasis. Brain 144:3036–3049. DOI: 10.1093/brain/awab206

46. Seaman MNJ. 2004. Cargo-selective endosomal sorting for retrieval to the Golgi requires retromer. The Journal of Cell Biology 165:111–122. DOI: 10.1083/jcb.200312034

47. Seaman MNJ, Michael McCaffery J, Emr SD. 1998. A Membrane Coat Complex Essential for Endosome-to-Golgi Retrograde Transport in Yeast. The Journal of Cell Biology 142:665–681. DOI: 10.1083/jcb.142.3.665

48. Simonetti B, Daly JL, Cullen PJ. 2023. Out of the ESCPE room: Emerging roles of endosomal SNX-BARS in receptor transport and host–pathogen interaction. Traffic 24:234–250. DOI: 10.1111/tra.12885

49. Simonetti B, Danson CM, Heesom KJ, Cullen PJ. 2017. Sequence-dependent cargo recognition by SNX-BARs mediates retromer-independent transport of CI-MPR. Journal of Cell Biology 216:3695–3712. DOI: 10.1083/jcb.201703015

50. Siniossoglou S, Pelham HRB. 2002. Vps51p Links the VFT Complex to the SNARE Tlg1p. Journal of Biological Chemistry 277:48318–48324. DOI: 10.1074/jbc.M209428200

51. Sleat DE, Lobel P. 1997. Ligand Binding Specificities of the Two Mannose 6-Phosphate Receptors. Journal of Biological Chemistry 272:731–738. DOI: 10.1074/jbc.272.2.731

52. Sleat DE, Wang Y, Sohar I, Lackland H, Li Y, Li H, Zheng H, Lobel P. 2006. Identification and Validation of Mannose 6-Phosphate Glycoproteins in Human Plasma Reveal a Wide Range of Lysosomal and Non-lysosomal Proteins. Molecular C Cellular Proteomics 5:1942–1956. DOI: 10.1074/mcp.M600030-MCP200

53. Söllner T, Bennett MK, Whiteheart SW, Scheller RH, Rothman JE. 1993. A protein assembly-disassembly pathway in vitro that may correspond to sequential steps of synaptic vesicle docking, activation, and fusion. Cell 75:409–418. DOI: 10.1016/0092-8674(93)90376-2

54. Tharkeshwar AK, Demedts D, Annaert W. 2020. Superparamagnetic Nanoparticles for Lysosome Isolation to Identify Spatial Alterations in Lysosomal Protein and Lipid Composition. STAR Protocols 1:100122. DOI: 10.1016/j.xpro.2020.100122

55. Toshima JY, Toshima J. 2024. Transport mechanisms between the endocytic, recycling, and biosynthetic pathways via endosomes and the trans-Golgi network. Frontiers in Cell and Developmental Biology 12:1464337. DOI: 10.3389/fcell.2024.1464337

56. Tyanova S, Temu T, Sinitcyn P, Carlson A, Hein MY, Geiger T, Mann M, Cox J. 2016. The Perseus computational platform for comprehensive analysis of (prote)omics data. Nature Methods 13:731–740. DOI: 10.1038/nmeth.3901

57. Wei J, Zhang Y-Y, Luo J, Wang J-Q, Zhou Y-X, Miao H-H, Shi X-J, Qu Y-X, Xu J, Li B-L, Song B-L. 2017. The GARP Complex Is Involved in Intracellular Cholesterol Transport via Targeting NPC2 to Lysosomes. Cell Reports 19:2823–2835. DOI: 10.1016/j.celrep.2017.06.012

58. Wenzel EM, Schultz SW, Schink KO, Pedersen NM, Nähse V, Carlson A, Brech A, Stenmark H, Raiborg C. 2018. Concerted ESCRT and clathrin recruitment waves define the timing and morphology of intraluminal vesicle formation. Nature Communications 9:2932. DOI: 10.1038/s41467-018-05345-8

59. Wickham H. 2016. Data Analysis. In: Wickham H (Ed). Ggplot2: Elegant Graphics for Data Analysis. Springer International Publishing. p. 189–201. DOI: 10.1007/978-3-319-24277-4_9

60. Willenborg M, Schmidt CK, Braun P, Landgrebe J, Von Figura K, Saftig P, Eskelinen E-L. 2005. Mannose 6-phosphate receptors, Niemann-Pick C2 protein, and lysosomal cholesterol accumulation. Journal of Lipid Research 46:2559–2569. DOI: 10.1194/jlr.M500131-JLR200

61. Wollert T, Hurley JH. 2010. Molecular mechanism of multivesicular body biogenesis by ESCRT complexes. Nature 464:864–869. DOI: 10.1038/nature08849

62. Yu G, Wang L-G, Han Y, He Q-Y. 2012. clusterProfiler: an R Package for Comparing Biological Themes Among Gene Clusters. OMICS: A Journal of Integrative Biology 16:284–287. DOI: 10.1089/omi.2011.0118

