## supplementary Figures for "Systematic Analysis of GARP- and EARP-Mediated Trafficking Pathways Using Lysosomal Proteomics"

**Supplementary Figures Sivalingam et al., 2026**

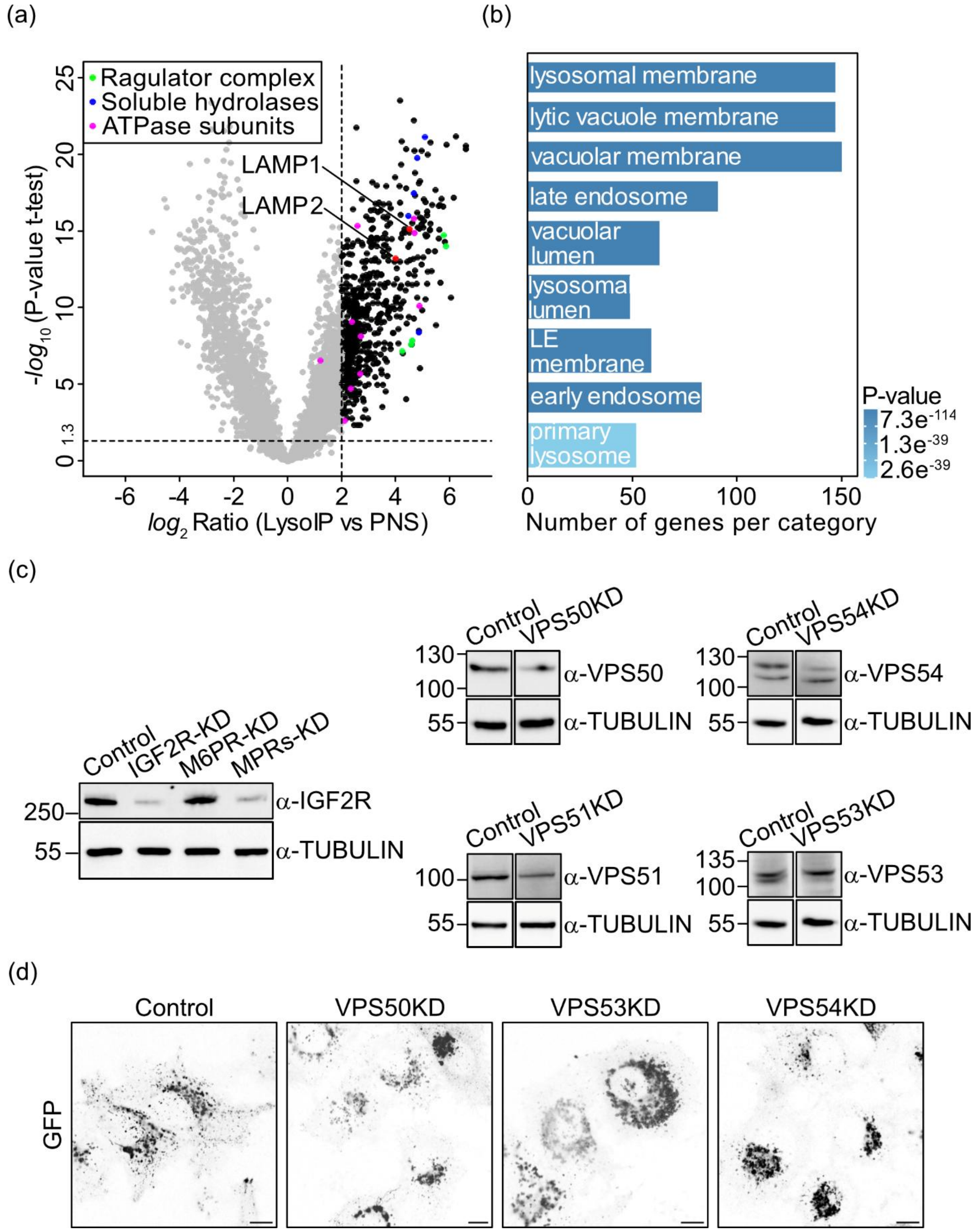

Fig. S1: **Characterization of Lyso-IP enrichment and validation of knockdown efficiency.** (a) Volcano plot showing proteins enriched in Lyso-IP from cells expressing LAMP1-GFP compared to the post-nuclear supernatant (PNS) ($log_2$ Ratio Lyso-IP > 2, -$log_{10}$ P-value > 1.3, n=15), with LAMP1 and LAMP2 serving as lysosomal markers (red). Furthermore, ATPase subunits including VA0D1, VATG1, VATB2, VATC1, VATE1, VAS1, VATF, VPP1, VATH, and VATD are highlighted in magenta, Ragulator complex subunits (LTOR1-5) in green, and soluble lysosomal hydrolases such as CATD, CATB, CATZ, HEXA, HEXB, and BGLR in blue. (b) GO term analysis highlighting the lysosomal-associated enrichment. (c) Western blots showing siRNA-mediated knockdown efficiencies: in the double knockdown of M6PR and IGF2R, we saw a 94 % reduction of Igf2r levels, 42 % for VPS50 (EARP-specific), 45 % for VPS54 (GARP-specific), 73 % for VPS51, and 50 % for VPS53, with tubulin used as a loading control. (d) Live-cell images of control, VPS50, VPS53, and VPS54 knockdown (KD) cells highlighting their distinct phenotypes. *Scale bar: 10 µm*.

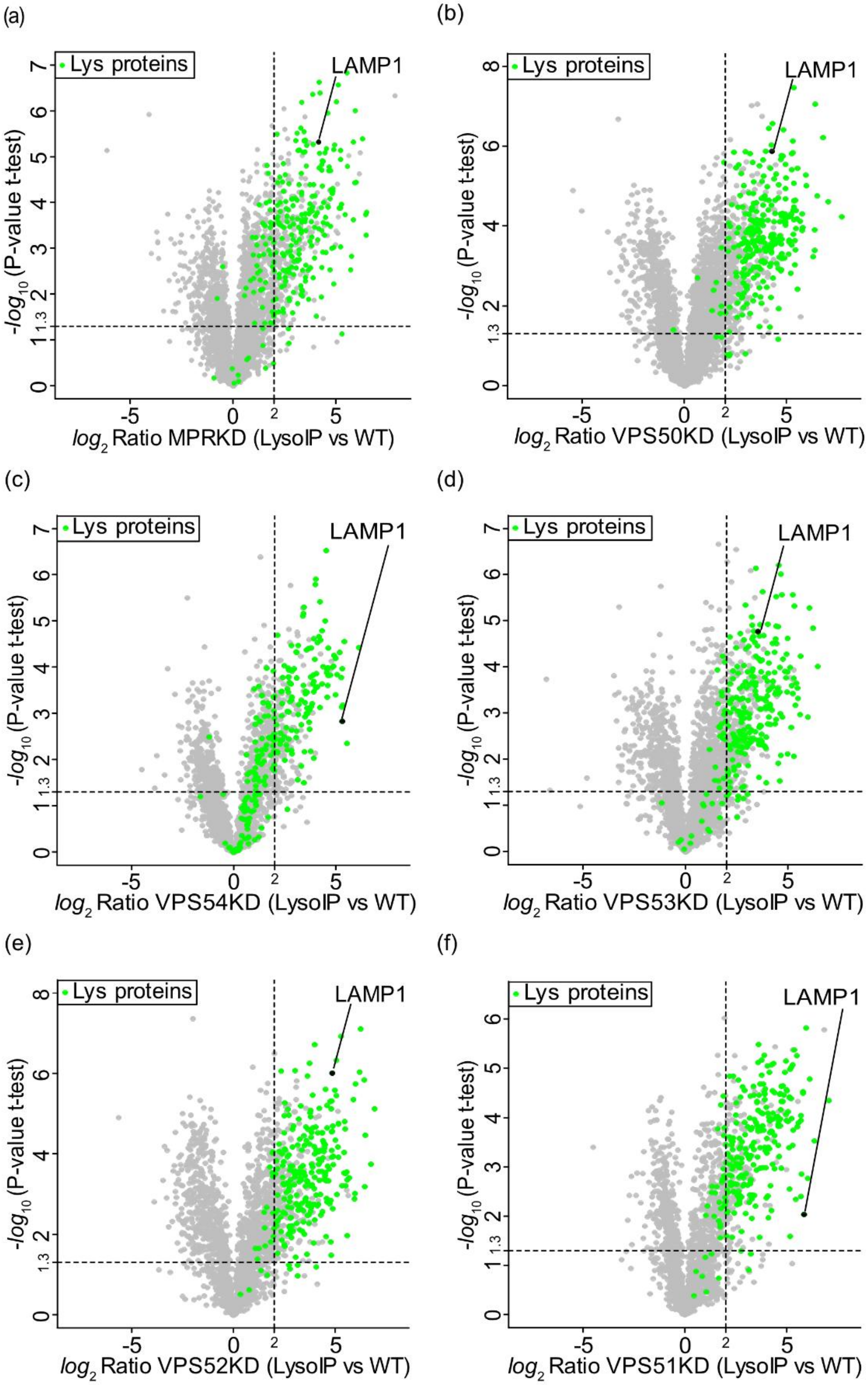


Sivalingam et al. 2026, Figure S2

Fig. S2: **Effect of MPR and GARP/EARP knockdowns on lysosomal protein enrichment.** Volcano plots showing Lyso-IP enrichment from (a) cation-dependent and -independent MPR knockdown, (b) VPS50 (EARP-specific) knockdown, (c) VPS54 (GARP-specific) knockdown, (d) VPS53 knockdown, (e) VPS52 knockdown, and (f) VPS51 knockdown. Previously defined lysosomal proteins (n = 265) are highlighted in green, illustrating differential enrichment patterns across knockdowns. Among the 265 lysosomal proteins, 62 lost lysosomal in MPR depleted lysosomes, 16 upon VPS50 depletion, 37 following VPS51 knockdown, 25 in VPS52 depletion, 43 upon VPS53 knockdown, and 119 lysosomal proteins following VPS54 depletion (n = 3).

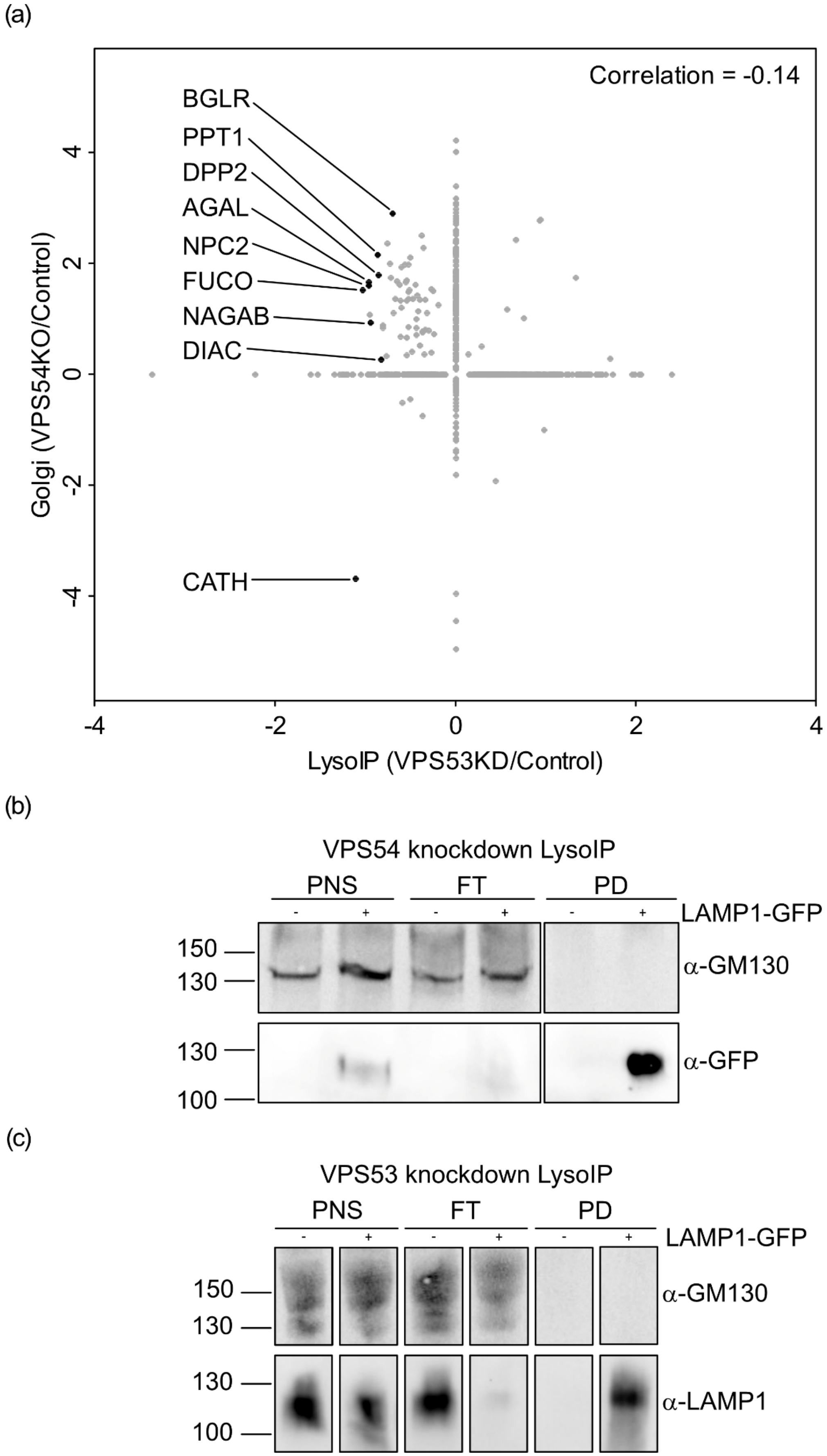

(a)
Correlation = -0.14
BGLR
PPT1
DPP2
AGAL
NPC2
FUCO
NAGAB
DIAC
CATH
4
2
0
-2
-4
Golgi (VPS54KO/Control)
-4
-2
0
2
4
LysoIP (VPS53KD/Control)
(b)
VPS54 knockdown LysoIP
PNS
FT
PD
-
+
-
+
-
+
LAMP1-GFP
150
130
α-GM130
130
100
α-GFP
(c)
VPS53 knockdown LysoIP
PNS
FT
PD
-
+
-
+
-
+
LAMP1-GFP
150
130
α-GM130
130
100
α-LAMP1

Fig. S3: **Golgi retention of lysosomal enzymes upon GARP complex knockdown.** (a) Correlation analysis between the Lyso-IP from VPS53 knockdown cells and the Golgi-enriched fraction from VPS54 knockout (wobbler) mice (Petit *et al.*, 2020) highlights a set of lysosomal enzymes, including PPT1, NAGAB, NPC2, glucuronidase (BGLR), peptidase DPP2, galactosidase (AGAL), fucosidase FUCO, and acetylchitobiase DIAC, that are de-enriched in Lyso-IP in VPS53KD but enriched in the Golgi fraction of VPS54Ko mice, indicating their Golgi retention upon GARP loss. (b) Western blot of Lyso-IP from VPS54 knockdown cells with and without LAMP1-GFP expression, probed with antibodies against the Golgi marker GM130 and GFP. Samples include 2.5 % (v/v) of the post-nuclear supernatant (PNS) and flow-through (FT), and 12.5 % (v/v) of the pulldown (PD). The absence of the GM130 signal confirms that Golgi contamination is negligible in the Lyso-IP fraction. (c) Western blot of Lyso-IP from VPS53 knockdown cells with and without LAMP1-GFP expression, probed with GM130 and LAMP1 antibodies, further confirming the absence of Golgi contamination in the Lyso-IP.

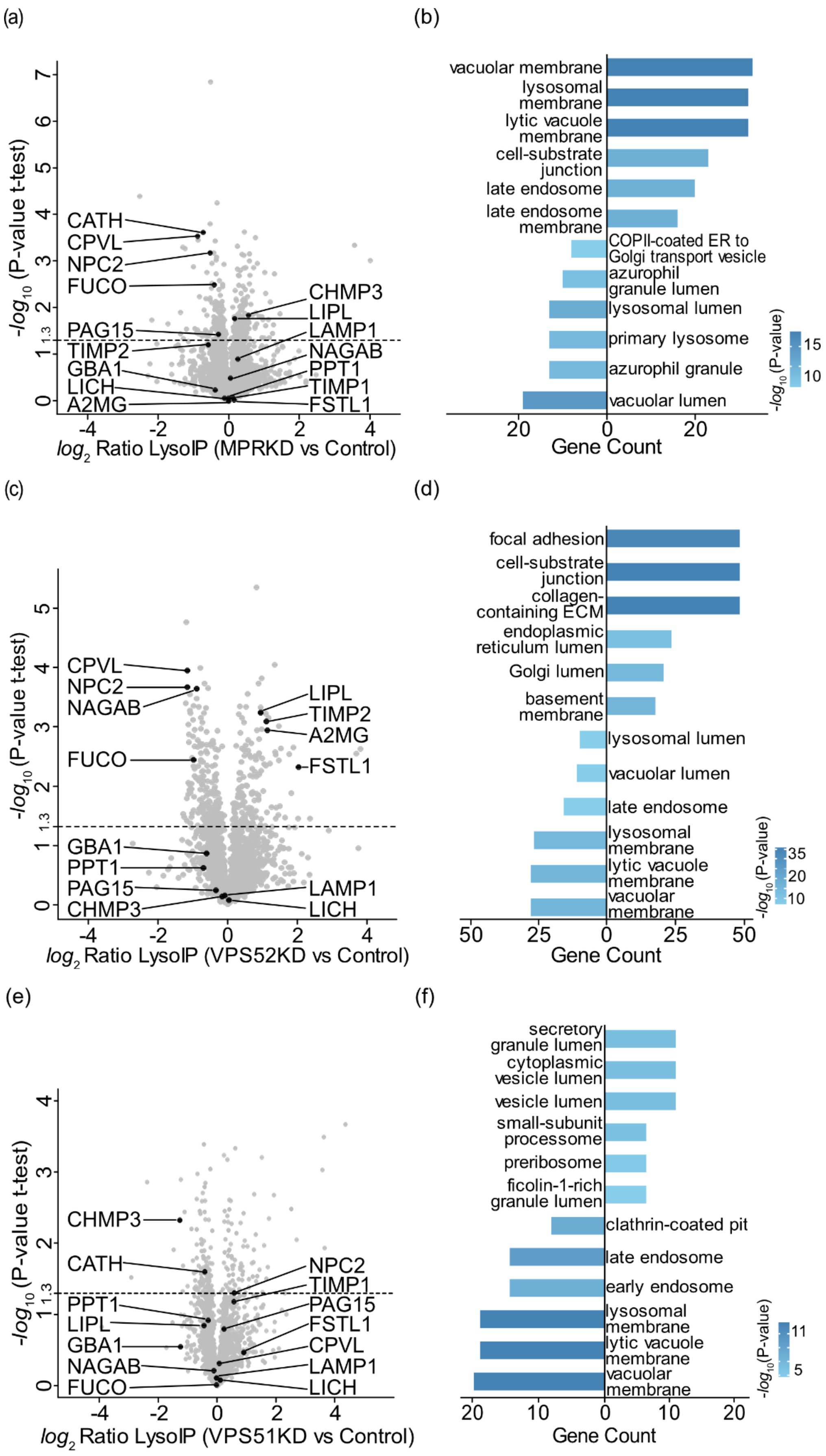

(a)
(b)
-log10 (P-value t-test)
log2 Ratio LysoIP (MPRKD vs Control)
CATH
CPVL
NPC2
FUCO
PAG15
TIMP2
GBA1
LICH
A2MG
CHMP3
LIPL
LAMP1
NAGAB
PPT1
TIMP1
FSTL1
vacuolar membrane
lysosomal membrane
lytic vacuole membrane
cell-substrate junction
late endosome
late endosome membrane
COPII-coated ER to Golgi transport vesicle
azurophil granule lumen
lysosomal lumen
primary lysosome
azurophil granule
vacuolar lumen
Gene Count
-log10(P-value)
(c)
(d)
log2 Ratio LysoIP (VPS52KD vs Control)
CPVL
NPC2
NAGAB
FUCO
GBA1
PPT1
PAG15
CHMP3
LIPL
TIMP2
A2MG
FSTL1
LAMP1
LICH
focal adhesion
cell-substrate junction
collagen-containing ECM
endoplasmic reticulum lumen
Golgi lumen
basement membrane
lysosomal lumen
vacuolar lumen
late endosome
lysosomal membrane
lytic vacuole membrane
vacuolar membrane
(e)
(f)
log2 Ratio LysoIP (VPS51KD vs Control)
CHMP3
CATH
PPT1
LIPL
GBA1
NAGAB
FUCO
NPC2
TIMP1
PAG15
FSTL1
CPVL
LAMP1
LICH
secretory granule lumen
cytoplasmic vesicle lumen
vesicle lumen
small-subunit processome
preribosome
ficolin-1-rich granule lumen
clathrin-coated pit
late endosome
early endosome
lysosomal membrane
lytic vacuole membrane
vacuolar membrane

Fig. S4: **Variations in lysosomal proteomes upon MPR and GARP/EARP knockdowns.** (a) Volcano plot of Lyso-IP compared to a control Lyso-IP from cation-dependent and -independent MPR knockdown. Lysosomal enzymes such as carboxypeptidase CPVL, the fucosidase FUCO, pro-cathepsin H (CATH), lysosomal cholesterol transporter NPC2, and phospholipase PAG15 are de-enriched in the MPR depleted lysosomes. Whereas, enrichment of the ESCRT-III component CHMP3 and lipoprotein lipase (LIPL) are observed. No significant changes in the levels of lysosomal acid lipase (LIPA), glucosylceramidase (GBA1), follistatin-related protein 1 (FSTL1), metalloproteinase inhibitors TIMP1 and TIMP2, alpha-2-macroglobulin (A2MG), palmitoyl-protein thioesterase 1 (PPT1), or alpha-N-acetylgalactosaminidase (NAGAB) were detected, in contrast to the changes observed upon GARP and EARP depletion. (b) GO term analysis of significantly enriched (right) and de-enriched (left) proteins (-$log_{10}$ P-value > 1.3). (c) Volcano plot of VPS52 knockdown Lyso-IP compared to a control Lyso-IP showed similar observations as in VPS53KD Lyso-IP including the de-enrichment of lysosomal enzymes (CPVL, NPC2, NAGAB, and FUCO) and the enrichment of secreted proteins such as LIPL, TIMP2, A2MG, and FSTL1. The levels of CHMP3, PPT1, LICH, GBA1, and PAG15 remain unchanged. (d) GO term analysis of significantly enriched and de-enriched proteins from VPS52 knockdown reveals impaired lysosomal enzyme targeting and enhanced cargo retention. (e) Volcano plot of VPS51 knockdown Lyso-IP compared to a control Lyso-IP highlights the de-enrichment of CHMP3 and CATH, and insignificant changes of PPT1, LIPL, GBA1, NAGAB, FUCO, LICH, CPVL, FSTL1, PAG15, TIMP1, and NPC2. (f) Corresponding GO term enrichment analysis of significantly enriched and de-enriched proteins.

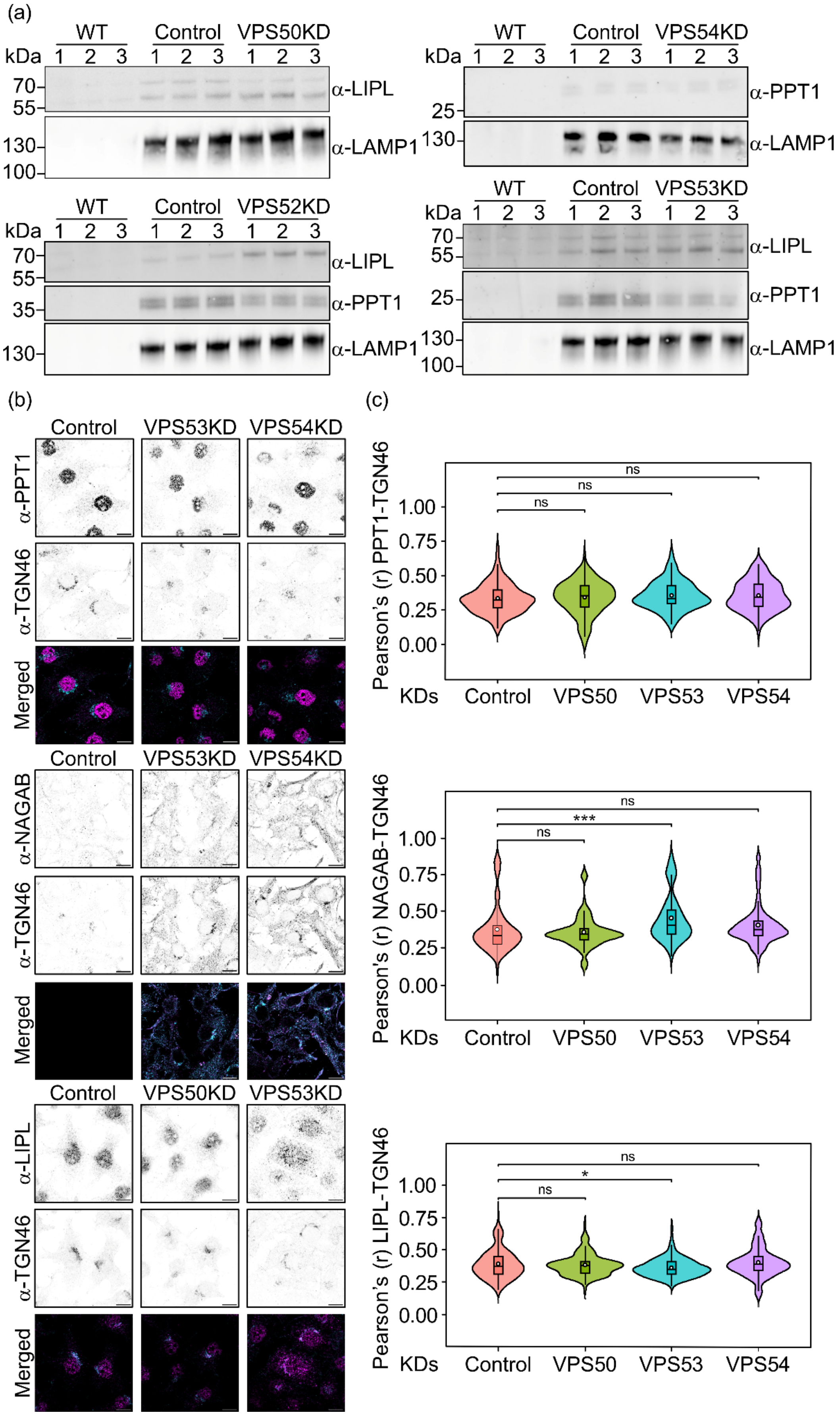

(a)
WT
Control
VPS50KD
kDa
1 2 3 1 2 3 1 2 3
70
55
α-LIPL
130
100
α-LAMP1
WT
Control
VPS54KD
kDa
1 2 3 1 2 3 1 2 3
25
α-PPT1
130
α-LAMP1
WT
Control
VPS52KD
kDa
1 2 3 1 2 3 1 2 3
70
55
α-LIPL
35
α-PPT1
130
α-LAMP1
WT
Control
VPS53KD
kDa
1 2 3 1 2 3 1 2 3
70
55
α-LIPL
25
α-PPT1
130
100
α-LAMP1
(b)
Control
VPS53KD
VPS54KD
α-PPT1
α-TGN46
Merged
Control
VPS53KD
VPS54KD
α-NAGAB
α-TGN46
Merged
Control
VPS50KD
VPS53KD
α-LIPL
α-TGN46
Merged
(c)
Pearson's (r) PPT1-TGN46
1.00
0.75
0.50
0.25
0.00
ns
ns
ns
KDs
Control
VPS50
VPS53
VPS54
Pearson's (r) NAGAB-TGN46
1.00
0.75
0.50
0.25
0.00
ns
***
ns
KDs
Control
VPS50
VPS53
VPS54
Pearson's (r) LIPL-TGN46
1.00
0.75
0.50
0.25
0.00
ns
*
ns
KDs
Control
VPS50
VPS53
VPS54

Fig. S5: **Effects of retrograde and recycling pathways on selected proteins.** (a) Western blot showing Lyso-IP samples from HeLa-WT, LAMP1-GFP, and LAMP1-GFP cells with VPS50, VPS54, VPS52, or VPS53 knockdown (n = 3). Blots were probed with antibodies against LIPL, PPT1, and LAMP1 to quantify and validate the proposed effects. (b) Immunostaining of PPT1, NAGAB, and LIPL with TGN46 in control and GARP/EARP knockdown cells. *Scale bar: 10 µm*. (c) Colocalization quantification of b represented as violin plots, with significance indicated by stars (*$p < 0.05$, **$p < 0.01$, ***$p < 0.001$, ****$p < 0.0001$). ($n = 3$, with $n \geq 104$ cells quantified per replicate.
